# Rare tRNAs with A-starting anticodons are functional and show codon-box dependent fitness effects

**DOI:** 10.64898/2026.05.15.725136

**Authors:** Parth K Raval, Sungbin Lim, Jenna Gallie, Deepa Agashe

## Abstract

Across the tree of life, there is immense diversity in transfer RNA (tRNA) gene copy number, anticodons, backbone sequences, and post-transcriptional modifications. Curiously, tRNAs with adenine-starting anticodons (tRNA_ANN_) are either absent or post-transcriptionally modified, suggesting purifying selection on unmodified tRNA_ANN_, e.g., due to non-functionality or toxicity. However, the functionality and fitness effects of ANN anticodons in their native tRNA backbones have not been measured. We generated tRNAs carrying all theoretically possible ANN anticodons, expressed them in *Escherichia coli* (genomically or on plasmids), tested functionality, and assessed fitness effects in various growth conditions. Contrary to expectation, every tRNA_ANN_ expressed from the genome was folded, matured, translationally active and substituted for native tRNA without severe fitness costs. Overexpression from a plasmid was also neutral in 50% of tRNA_ANN_-media combinations tested. In the remaining 50% of cases, tRNA_ANN_ decoding four-fold degenerate codon boxes (4D tRNA_ANN_) often improved fitness, whereas those decoding two-fold degenerate codon boxes (2D tRNA_ANN_) tended to impair fitness. We speculate that increased probability of miscoding by 2D tRNA_ANN_ may explain why they are more costly than 4D tRNA_ANN_, and hope that future work will elucidate the underlying mechanisms. Our results demonstrate a lack of strong and pervasive fitness costs of tRNA_ANN_ suggesting overall weak purifying selection, and sharpen the paradox of the near-universal absence of unmodified tRNA_ANN_.

## INTRODUCTION

Transfer RNA (tRNA) pools, largely determined by tRNA gene copy numbers (GCN), play a crucial role in maintaining translation rate, translation accuracy, growth rate and overall fitness (Young et al. 1976; Dong et al. 1996; Scott et al. 2010; Scott and Hwa 2011; Klumpp et al. 2013; Wilusz 2015; Hu and Lercher 2021; Raval et al. 2023). Within species, tRNA repertoires adapt in response to various environmental factors (Nilsson et al. 2006; Iben et al. 2011; Bedhomme et al. 2019; Ayan et al. 2020; Khomarbaghi et al. 2024), and across species, they echo the diversity of ecological niches (Goodenbour and Pan 2006; Vieira-Silva and Rocha 2010; Fujishima and Kanai 2014; Chan and Lowe 2016; Rak et al. 2018). For a given amino acid, the GCN of tRNAs with distinct anticodons for the same amino acid (‘tRNA isoacceptors’) varies across species. The GCN is often correlated with relative codon usage (Ikemura 1985; Dong et al. 1996; Berg and Kurland 1997; Elf et al. 2003; Rocha 2004; Novoa and Ribas de Pouplana 2012; McDonald et al. 2015), contributing to the diversity of tRNA repertoires. The effective cytosolic tRNA pools can also vary with expression levels from each tRNA gene copy (Kanaya et al. 1999; Dittmar et al. 2004; Dittmar et al. 2006; Bloom-Ackermann et al. 2014; Sagi et al. 2016; Raval et al. 2023) and differential charging across isoacceptors (Elf et al. 2003; Dittmar et al. 2005). Another layer of variability is generated by tRNA modifying enzymes (MEs) that can post-transcriptionally modify tRNAs at about a dozen different sites, particularly in the anticodon loop which extends the initially proposed wobble pairing (Crick 1966) to an expanding list of modified wobbles (Agris et al. 2007; Grosjean et al. 2010; Iben and Maraia 2012; Novoa et al. 2012; Agris et al. 2017; Maraia and Arimbasseri 2017; Agris et al. 2018; Suzuki 2021). Despite such numerous mechanisms that generate an immense diversity of functional tRNAs within and across species, tRNAs with specific anticodons are curiously either missing or rare (Maraia and Arimbasseri 2017; Diwan and Agashe 2018; Rak et al. 2018; Torres 2019; Lei and Burton 2020; Ehrlich et al. 2021; Pernod et al. 2021).

Of these seemingly ‘prohibited’ tRNAs, perhaps most striking is the lack of those carrying an unmodified adenine at the 34^th^ (wobble) position (henceforth, tRNA_ANN_). A total of eight ANN anticodons are theoretically possible for the eight family-codon boxes with four-fold degeneracy (‘4D’; all four codons within a box, differing only at the 3^rd^ base, encode the same amino acid), and another eight for the split-codon boxes with two-fold degeneracy (‘2D’; the four codons within a box encode different amino acids) (Fig. 1A). tRNA_ANN_ carrying each of these theoretically possible ANN anticodons are a single point mutation away from tRNA_UNN_, tRNA_GNN_, and tRNA_CNN_ (collectively, ‘tRNA_BNN_’) in the same codon box, and should occur frequently in bacterial populations. For instance, with a mutation rate of 9.1×10^−11^ per base per generation, an *Escherichia coli* population that grows from ca. 1,000 cells to 1 billion cells (e.g., a 1 mL culture grown in rich medium overnight) will sample at least one mutation leading to a tRNA_ANN_. However, seven 2D codon tRNA_ANN_ and one 4D codon tRNA_ANN_ are universally absent (Fig. 1A). In addition, seven more 4D tRNA_ANN_ are absent in prokaryotes, though Leu-tRNA_AAG_ and Thr-tRNA_AGT_ do occur in some bacteria (Andachi et al. 1987; Borén et al. 1993; Inagaki et al. 1995; Phillips and de Crécy-Lagard 2011; Diwan and Agashe 2018; Ehrlich et al. 2021). Curiously, these latter seven 4D codon box tRNA_ANN_ are also absent from eukaryotic organelles of bacterial origin and from bacteriophages (Hatfull 2015; Pope et al. 2015; Morgado and Vicente 2019; Ehrlich et al. 2021). Hence, comparative genomics alone underscore the persistent rarity of tRNA_ANN_, suggesting that their expression may not be well-tolerated.

**Figure 1:**
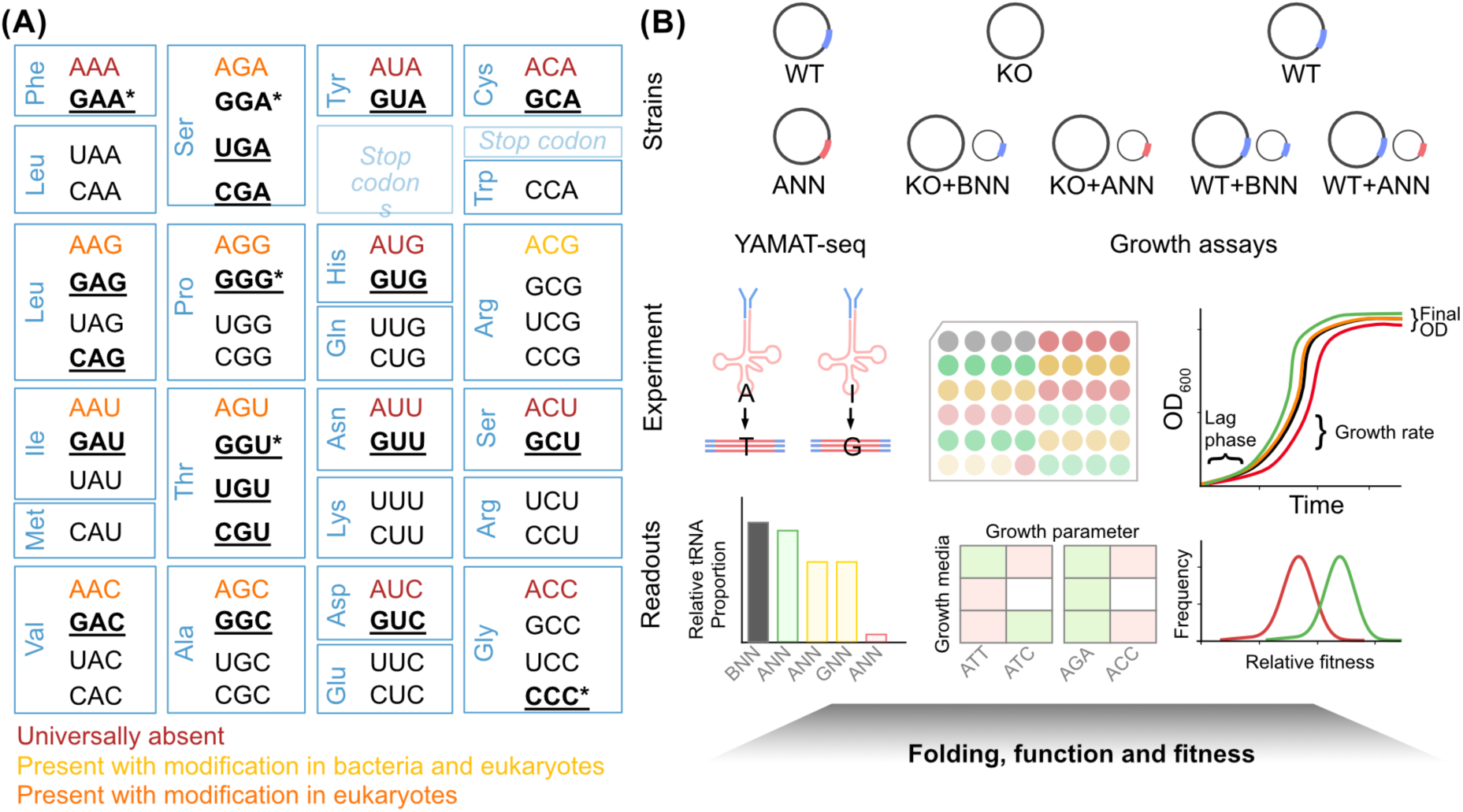
Theoretically possible tRNA_ANN_, their potential cellular consequences, and our approach to test their fitness effects. **(A)** All theoretically possible anticodons (written from 5’-to-3’) organized across 4D and 2D codon boxes in the codon table. ANN anticodons that are universally rare and modified in different domains (based on (Ehrlich et al. 2021)) are color coded as per the key on the bottom. Anticodons of the native tRNA_BNN_ that were mutated to tRNA_ANN_ in this study (within their respective codon boxes) are indicated in bold, of which, BNN anticodons whose tRNA_ANN_ variants were expressed from a plasmid are underlined, and those expressed from the genome are indicated by asterisks. **(B)** To test whether tRNA_ANN_ are expressed, folded and matured, we utilized YAMAT-seq (Y-shaped Adapter-ligated MAture TRNA sequencing) (Shigematsu et al. 2017; Ayan et al. 2020; Raval et al. 2023). To test for folding, functionality and fitness we used the wild type *E. coli* MG1655 (WT), strains with B34A substitution on the genome (‘ANN strains’), strains lacking native tRNA genes (knockouts or ‘KOs’), KOs complemented by tRNA_ANN_ or tRNA_BNN_ genes from a plasmid, and finally WT expressing tRNA_ANN_ or tRNA_BNN_ from a plasmid. The functionality of tRNA_ANN_ was tested via their ability to substitute for the native tRNA_BNN_, as inferred by three growth parameters (lag phase, rate and final OD) estimated from OD_600_ vs time curves. Equivalent function would be inferred if WT and ANN strains have similar growth; if WT+ tRNA_ANN_ and WT+ tRNA_BNN_ have similar growth; and the poor growth of KOs is rescued equally well by plasmid-expressed tRNA_ANN_ and tRNA_BNN_. For WT+ tRNA_ANN_ and WT+ tRNA_BNN_, the magnitude of fitness effects of tRNA_ANN_ was measured as the relative change in growth parameters for each of these comparisons.

There are some exceptions where tRNA_ANN_ are genomically encoded; however, their post-transcriptional processing again indicates poor tolerance of unmodified tRNA_ANN_. Both bacteria and eukaryotes encode Arg-tRNA_ACG_, which is in fact the preferred Arg-tRNA isoacceptor. But in this case, the A34 is post-transcriptionally masked via deamination by the modifying enzyme TadA in prokaryotes (Karcher and Bock 2009; Rafels-Ybern et al. 2019; Ehrlich et al. 2021) or ADAT in eukaryotes, resulting in tRNAs with inosine-34 (tRNA_ICG_) in the cytosol (Gerber and Keller 1999; Wolf et al. 2002; Torres et al. 2014) and in the chloroplast stroma (Delannoy et al. 2009; Karcher and Bock 2009). Eukaryotes also encode and similarly modify six additional 4D codon box tRNA_ANN_ (Torres et al. 2014; Rafels-Ybern et al. 2018) (Fig. 1A). The only remaining 4D codon box tRNA_ANN_ (Gly-tRNA_ACC_) is universally absent, likely because it is a poor substrate for ADAT (Saint-Léger et al. 2016) and TadA (Borén et al. 1993). The A-to-I modification appears to be essential for efficient translation in bacteria (Wolf et al. 2002), eukaryotes (Torres et al. 2021) and chloroplasts (Delannoy et al. 2009). Notably, both tRNA_ANN_ and TadA/ADAT seem to be absent in archaea and mitochondria (Ehrlich et al. 2021). Together, the strict co-occurrence of tRNA_ANN_ and TadA/ADAT and the universal lack of the poor TadA/ADAT substrate tRNA_ANN_ point to ancient and strong purifying selection against unmodified tRNA_ANN_, which has likely persisted through the four billion years of life on Earth.

As noted above, single base mutations should frequently generate tRNA_ANN_ from tRNA_BNN_ genes, but are presumably removed by selection, e.g., due to their non-functionality or toxicity. However, *in vivo* tests of functionality and potential fitness impacts of unmodified tRNA_ANN_ are limited. Previous studies have largely focused on a few ANN anticodons in their native backbones or expressed multiple ANN from a single backbone, limiting the ability to generalize. For instance, Gly-tRNA_ACC_ is unmodified and functional in *E. coli* (Borén et al. 1993) and humans (Saint-Léger et al. 2016) and Pro-tRNA_AGG_ does not affect fitness or translational elongation rates in *Salmonella* (Chen et al. 2002). Both these tRNA_ANN_ were generated from the respective native tRNA_BNN_ backbones. In another study, all sixteen possible ANN anticodons were generated from a single *Methanocaldococcus jannaschii* Tyr-tRNA backbone, and these introduced tRNA_ANN_ decoded 1-20% of NNU codons in *E. coli* (Biddle et al. 2016; Schmitt et al. 2018; Schmitt et al. 2024). These observations suggest that unmodified tRNA_ANN_ are likely to be translationally active and non-lethal. However, the decoding capacities and modifications of tRNAs (including A-to-I) depend on the backbone sequence (Li et al. 1997; Qian et al. 1997; Nakanishi et al. 2005; Schmeing et al. 2011; Saint-Léger et al. 2016; Roura Frigolé et al. 2019); and fitness impacts of tRNAs depend on the ecological niche (Bloom-Ackermann et al. 2014; Li et al. 2016; Li and Zhang 2018; Gabzi et al. 2022; Raval et al. 2023). Therefore, validating function in native tRNA backbones and conducting systematic fitness assays across different growth media are both essential to quantify the hypothesized selection against tRNA_ANN_.

To this end, we introduced 20 tRNA_ANN_ via point mutation in the native tRNA genes of *E. coli*, and tested their expression, maturation and functionality. Our results provide some of the first systematic empirical evidence for folding, maturation and translational activity of a broad set of tRNA_ANN_ in an endogenous context. Most of the tRNA_ANN_ were accommodated in the cytosol unmodified, did not alter fitness under tested conditions, and a subset of tested tRNA_ANN_ evidently effectively replaced native tRNA_BNN_. However, within the tRNA_ANN_ that affected fitness, 2D and 4D tRNA_ANN_ differed from each other. 2D tRNA_ANN_ appeared prone to modification and sometimes reduced fitness when overexpressed, whereas 4D tRNA_ANN_ appeared unmodified and often improved fitness, especially in nutrient poor media. Our results show that tRNA_ANN_ are largely functional and in several cases even beneficial, deepening the mystery of the near-universal absence.

## METHODS

### Generating strains

We conducted all tRNA gene manipulations in *E. coli* MG1655 (the wild type, WT). The native tRNA genes were amplified from the genome and cloned into an IPTG-inducible low copy number plasmid (pACDH, a derivative of pACD with an ACYC origin of replication (Mangroo and RajBhandary 1995; Rao and Varchney 2001)) or a high copy number plasmid (pUC19 (Norrander et al. 1983)). We introduced an B34-to-A34 point mutation (‘B34A’) in the native tRNA_BNN_ genes via PCR mutagenesis (Hoa et al. 1989), followed by Sanger sequencing to confirm the mutation. To introduce tRNA_ANN_ on the genome, we utilized a scar-free, two-step allelic exchange method that utilizes the pKOV vector that carries a chloramphenicol resistance marker (Link et al. 1997). Briefly, we amplified the locus with the native tRNA gene of interest (along with ca. 200-300bp upstream and downstream regions) from the genome while also introducing a B34A substitution in the amplicon through PCR mutagenesis. We cloned the amplicons with mutated alleles into pKOV and transformed the WT with pKOV-tRNA_ANN_. We selected colonies with genomic integration of pKOV-tRNA_ANN_ on chloramphenicol (20 µg/ml) at 43°C (where the pKOV plasmid replicates very inefficiently) and confirmed the chromosomal insertion via PCR. We selected cells that had removed pKOV, by growing the population first overnight in LB (Lysogeny Broth, Difco) with 5% sucrose at 37°C and then on LB agar with 5% sucrose to isolate single colonies at 37°C. The presence of sucrose in both these steps selects against cells with pKOV since the expression of *sacB* from pKOV is lethal. We attempted these experiments for thirteen genes (*serX*, *glyU*, *pheV*, *thrT*, *thrV*, *pheU*, *serW*, *asnV*, *aspT*, *thrW*, *serU*, *serT*, *leuU*) at least three times per gene; but only the first four yielded successful B34A mutations and were investigated further. *proL B34A* was introduced on the genome via the Datsenko-Wanner method (Datsenko and Wanner 2000). Briefly, kanamycin cassette was first inserted 60 bp downstream of *proL.* The *proL::Kan* region was amplified through a mutagenesis PCR that introduced G34A via the forward primer. The *proL G34A::Kan* amplicon was electroporated into the WT carrying plasmid pKD46 where the red recombinase switched the WT *proL* with *proL G34A::Kan*. The Kan cassette was cured using PCP20 resulting in the proL G34A strain. We screened the mutant strains for the intended mutation, and to rule out any other mutations, via PCR followed by Sanger sequencing and further confirmed them via Next Generation Sequencing (Illumina HiSeq PE150, 4-5 million reads per strain, >100x depth). We stored all the strains thus generated as glycerol cryostocks at –80°C.

### Measuring growth parameters

We revived strains from cryostocks by streaking onto LB (Lysogeny Broth, Difco) agar plates, with appropriate antibiotics where applicable. We inoculated individual colonies in LB and incubated at 37°C (180 rpm shaking) for 14-16 hours to grow the preculture. We set up growth measurement experiments by sub-culturing 1% v/v of precultures in 48 well microplates (Corning) in 500 µL growth medium: LB or M9 minimal medium (M9 salts, 1mM CaCl2, 2.5 mM MgSO4) supplemented with specific carbon and nitrogen sources (“GA”: 0.4% w/v glucose and cas amino acids; or carbon sources alone: glucose (Glu) 0.2% w/v, galactose (Gal) 0.2%, pyruvate (Pyr) 50 mM, or glycerol (Gly) 0.6% w/v) (Raval et al. 2023) and with appropriate antibiotics where applicable. To test for fitness costs that may accumulate over longer periods of time, we passaged the strains with a B34A substitution (six independent lines per strain) for eight days (about 80 generations) by diluting 1% v/v every 24h in 500 µL LB in 48 well microplates incubated at 37°C (180 rpm shaking). We stored cryostocks at fourth and eighth day, revived and measured their fitness as per above.

We estimated various growth proxies by measuring optical density at 600 nm (OD_600_) in a Tecan microplate reader, an automated system (LiconiX incubator, robotic arm and Tecan microplate reader) for strains where tRNA_ANN_ were introduced from the plasmid, or a microplate reader from Agilent Biotek (Epoch2) for strains with a B34A substitution. We included the WT in every microplate as a reference, to account for differences across plate readers and over time. We inferred growth rate by fitting an exponential equation to approximately the first one third of each growth curve (region where growth was exponential) using Curvefitter software (Delaney et al. 2013). We inferred the lag phase length as the time taken to reach early log phase, defined as approximately 1/3^rd^ of the final OD_600_ in a given growth medium (OD_600_ 0.2 for LB and M9 GA; 0.15 for M9 Glu, Gal and Gly; and 0.12 for M9 Pyr). We normalized fitness of WT carrying tRNA_ANN_, with that of WT carrying the respective tRNA_BNN_ control for each case of tRNA_ANN_ expression from a plasmid.

Finally, we approximated the overall fitness effect of each 2D codon tRNA_ANN_ expressed from high copy plasmid by qualitatively scoring its impact on the three growth parameters across six media, assigning +1, 0, and –1 to positive, neutral, and negative effects, respectively. We defined the sum of these scores, across media and across parameters, as the ‘fitness effect index’. We estimated the mistranslation likelihood for each tRNA_ANN_ as the ratio of non-cognate to cognate codon usage across the *E. coli* genome.

### Estimating adenine to inosine modification and mature tRNA levels

We measured the relative abundance of each tRNA isotype, for strains with B34A substitutions. Prior work suggests that adenine to inosine modification can be reliably estimated using the frequency of cytosine in the complementary strand, i.e., unmodified A34 appears as T34 in reverse-complemented cDNA, whereas I34 appears as C34 (Schmitt et al. 2024). We therefore utilized YAMAT-Seq (Shigematsu et al. 2017; Ayan et al. 2020) as optimized previously for *E. coli* (Raval et al. 2023). Briefly, we grew three biological replicates of each strain in LB till mid-log phase (ca. OD 0.3) and isolated total RNA. After deacylation, we ligated Y-shaped, DNA/RNA hybrid adapters (Shigematsu et al. 2017) (Eurofins) to the 5’-NCCA-3’ and 3’-inorganic phosphate-5’ ends of uncharged tRNAs, reverse transcribed, and further amplified the cDNA products by PCR with sample-specific indices (Illumina). After a quality and quantity check through bioanalyzer (Agilent DNA7500 Series II kit and Agilent 2100 Bioanalyzer Systems), we combined equimolar amounts of each sample and further purified the fully assembled cDNA (with adapters and indices ligated at both ends of the tRNA sequence) by separating them from unligated adapters based on size on a 5% poly-acrylamide gel. The final products were sequenced using an Illumina NextSeq 550 High-Output 2×75 bp kit (Single-end, 150 bp reads) at the Max Planck Institute for Evolutionary Biology. YAMAT-seq for proL-ANN (with WT control) was performed with size-based separation by magnetic beads (Ampure XP beads) followed by verification of the size of purified products (200-223 bp) using TapeStation DNA D1000 HS Screen Tape and sequencing on the Novaseq platform (paired end, 150 bp) at the National Centre for Biological Sciences.

We sorted the combined raw reads into reads derived from individual strains (available from NCBI GEO; accession number GSE328815) based on the unique Illumina barcodes. We downloaded all native tRNA sequences predicted by GtRNAdb 2.0 (Chan and Lowe 2009) for WT and manually added additional tRNA_ANN_ sequences by introducing B34A substitution in the sequence of tRNA_BNN_ isoacceptors mutated in our study (e.g. to get the gly-tRNA_ACC_ sequence, we replaced the C34 in gly-tRNA_CCC_ sequence with A34). We mapped bases 80-151 (expected to tRNA sequences) of the raw reads from each strain against the collection of native and ANN tRNA sequences using Geneious Prime (version 11.1.4). We used the following previously mapping criteria: 10 % mismatch/gap rate, max 2-bp gap size, max. 5 ambiguities; no iterations, discard reads that align equally well to multiple reference sequences (Khomarbaghi et al. 2024). On average, at least around 95% of the reads aligned with tRNAs, the number of reads aligned to each tRNA isoacceptor were exported by Geneious and used for calculating within-sample proportions of each tRNA isoacceptor.

### Analyses of tRNA sequences from GtRNA DB and TadA homologues

We downloaded all predicted tRNA sequences from all bacteria and archaea available on the Genomic tRNA database, GtRNAdb (Chan and Lowe 2016; Thornlow et al. 2020) including the taxonomy of each species. For 14 genera where at least one tRNA_ANN_ (other than Arg-tRNA_ACG_) was present in all of its species (a total of 273 species across genera) we attempted to retrieve protein coding sequences (CDS) as nucleotide sequences from NCBI Genome Datasets (v18.24.0), using the taxonomic IDs provided on GtRNAdb. We could retrieve genomes of 263 species using this approach for which we retrieved genome AT content from the metadata of the genome assemblies, and calculated codon usage from nucleotide CDS (eight Lactobacilli and two Streptococci species, the genomes could not be fetched readily using the taxonomy IDs). Lastly, out of 555 bacterial genomes from GtRNAdb with at least one tRNA_ANN_ (other than Arg-tRNA_ACG_), we could retrieve amino acid CDS for 539 genomes which were used to searched for TadA homologues using the *E. coli tadA* amino acid sequence as a query (diamond BLAST v2.1.24.178 (Buchfink et al. 2014); e-value<10e-6, percentage identity >30%, query coverage >50%).

### Data analysis

We used in-house python scripts (except for YAMAT-Seq) for all statistical analysis and generating plots. We used Mann–Whitney U tests to compare the fitness of strains expressing tRNA_ANN_ with relevant control strains (WT or strain expressing tRNA_BNN_). To quantify skew in fitness effects relative to zero (neutral), we computed a zero-anchored quantile asymmetry metric (Q-asym_0_) defined as Q(+)_0.9_ - |Q(-)_0.1_| / Q(+)_0.9_ + |Q(-)_0.1_|, where Q(+)_0.9_ is the 90^th^ percentile of positive values of log_2_(magnitude of fitness) and Q(-)_0.1_ the 10^th^ percentile of negative values. This metric compares the extent of the positive and negative tails relative to zero, yielding a normalized measure of tail imbalance scaled between −1 (deleterious skew) and +1 (beneficial skew), with 0 indicating symmetry. While similar in essence to Bowley’s coefficient (a measure based on quantiles and median), Q-asym_0_ is zero-anchored. Significance was assessed by bootstrap (10,000 resamples), with percentile-based 95% confidence intervals; skew was considered significant when the interval excluded zero. All raw data used for analysis are provided in the source data files for each figure, along with the statistics; in-house scripts to conduct the analyses and plots are available on Zenodo (10.5281/zenodo.23323378).

## RESULTS

### Generation of an *E. coli* strain library encompassing all theoretically possible A-starting anticodons

In theory, sixteen anticodons can start with an adenosine, but fifteen of these are absent from bacterial tRNA repertoires, including that of *E. coli* MG1655 (the wild type, WT). The WT tRNA repertoire consists of 37 genes for tRNAs with G-, U- or C-starting anticodons (‘tRNA_BNN_’) that decode family or four-fold degenerate codon boxes (‘4D’ codon boxes, where all four codons within a codon box are assigned to the same amino acid) and 45 genes for tRNAs that decode split or two-fold degenerate codon boxes (‘2D’ codon boxes, where the four codons within a codon box are split across two different amino acids). Thus, a total of 82 native tRNA_BNN_ genes (Fig. S1) could generate a tRNA with an A-starting anticodon (‘tRNA_ANN_’) through a single substitution leading to adenosine at the 34^th^ base (‘B34A substitution’) (Fig. 1A, Fig. S1).

To test the functionality and fitness effects of tRNA_ANN_, we constructed tRNA_ANN_ genes by introducing B34A substitutions in 20 diverse WT tRNA_BNN_ genes (Fig. 1, Fig. S1). From each of the eight 2D codon boxes, we derived one tRNA_ANN_ from the native tRNA_GNN_ (mimicking possible transition mutations). Of the eight 4D codon boxes, one is decoded by tRNA_ACG_ whereas the remaining seven are decoded by different combinations of tRNA_BNN_ isoacceptors with different backbones (Fig. S2). We introduced B34A substitutions in a subset of these tRNA_BNN_ isoacceptor genes. We generated one tRNA_ANN_ each from proline, valine, alanine and glycine 4D codon boxes. For serine, leucine and threonine 4D codon boxes, we generated tRNA_ANN_ from a subset of isoacceptors differing from each other in their backbone sequences by more than 30%, resulting in two tRNA_ANN_ from leucine codon boxes, and three from the threonine and serine codon box.

This set of 20 tRNA_ANN_ genes thus covered all fifteen ANN anticodons absent across bacteria, with serine, leucine and threonine ANN generated from more than one backbone. We could successfully integrate five of these tRNA_ANN_ into the WT genome by replacing one copy of the corresponding native tRNA_BNN_ with its tRNA_ANN_ variant (Fig. 1A), generating strains with tRNA_ANN_ expressed from the genome (individually referred with the gene name followed by ‘ANN’, e.g. pheV-ANN, and collectively referred to as ‘ANN strains’). Next, we used seven *E. coli* strains with a native tRNA_BNN_ gene deletion (Raval et al. 2023) (e.g. ΔpheV; “KO” in Fig. 1), and expressed the respective tRNA_ANN_ from a high-copy plasmid (Raval et al. 2023). In the WT, we additionally expressed five tRNA_ANN_ from a low-copy plasmid and 19 tRNA_ANN_ from a high-copy plasmid. The strains thus generated (Table S1, Fig. 1) provided a comprehensive system to probe specific aspects of tRNA_ANN_ functionality and fitness effects (Fig. 1B-C).

### tRNA_ANN_ are translationally active

A parsimonious mechanism by which tRNA_ANN_ can affect cellular fitness is through translation rate and/or accuracy, although evidence for their translational activity remains indirect and limited (Borén et al. 1993; Biddle et al. 2016; Schmitt et al. 2018; Schmitt et al. 2024). A key prerequisite for translational activity is accurate post-transcriptional folding and maturation of the tRNA transcripts. To quantify the relative abundance of folded and mature tRNA_ANN_ within the total tRNA pool, and to identify A34-I34 modification, we utilized YAMAT-seq (Y-shaped Adapter-ligated MAture TRNA sequencing) (Shigematsu et al. 2017; Ayan et al. 2020; Raval et al. 2023). In this method, the two ends of Y-shaped adapters are ligated to the conserved 3′-CCA and to 5’-OH, respectively, in the acceptor arms of fully folded and matured tRNAs. Adapter-ligated tRNAs are then reverse-transcribed, PCR-amplified using adapter-specific primers, and the resulting libraries are sequenced and analyzed using a standard small-RNA sequencing workflow. Thus, the method only detects mature tRNAs. We quantified the relative abundance of each tRNA species in all five strains where we replaced native tRNA_BNN_ genes by the respective tRNA_ANN_ variants (Fig. 1B). For four of these ANN strains, we also included strains lacking genes coding for the corresponding native tRNA_BNN_ (Raval et al. 2023), as a control for potential compensatory upregulation from the remaining copies of the deleted BNN gene.

Among the attempted experiments to replace a tRNA_BNN_ gene with a tRNA_ANN_ gene in the genome, we succeeded in generating one 2D and four 4D ANN strains. As expected, deleting the tRNA_BNN_ gene tended to decrease the amount of tRNA_BNN_, though this was statistically significant only for thrT and glyU (compare grey and orange bars in Fig. 2A). Likewise, when the tRNA_BNN_ gene carried a B34A substitution, the amount of tRNA_BNN_ was also lower than WT, except for the pheV-AAA strain (compare gray and first cyan bars in Fig. 2A). Importantly, in these ANN strains we also detected a substantial amount of tRNA_ANN_, indicating their successful expression and maturation (Fig. 2A). The only exception was the pheV-AAA strain, which produced little Phe-tRNA_AAA_ though it had a similar proportion of Phe-tRNA_GAA_ as WT (Fig. 2A). This suggested that at least a fraction of Phe-tRNA_GAA_ in pheV-AAA is due to A34-to-I34 modification of Phe-tRNA_AAA_, which would be detected in sequencing as tRNA_GAA_ (Fig. 1B). In contrast, all four 4D tRNA_ANN_ that we tested were expressed as mature tRNAs unmodified at the A34 position (henceforth simply ‘unmodified’, though modifications other than A34-to-I are likely present) (Fig. 2A). Finally, the total proportion of tRNA_BNN_ and tRNA_ANN_ produced by the ANN strains was consistently similar to the WT (compare the gray and last green bar, Fig. 2A). The observation that 4D tRNA_ANN_ proportions were similar to WT or added up to WT together with tRNA_BNN_ suggested that tRNA_ANN_ were tolerated unmodified in cells, without any inhibition of expression from *B34A* alleles or post-transcriptional degradation. Furthermore, in strains carrying tRNA_ANN_, relative levels of isoacceptors or other tRNA species were unaltered (Fig. S3), suggesting that tRNA_ANN_ expression is tolerated without major changes in relative cytosolic tRNA composition. Taken together, YAMAT-seq confirmed that both 4D and 2D codon tRNA_ANN_ are expressed, correctly folded, and fully mature akin to native tRNA_BNN_. The 4D codon tRNA_ANN_ remained unmodified, whereas the 2D codon tRNA_ANN_ was modified to tRNA_INN_.

**Figure 2:**
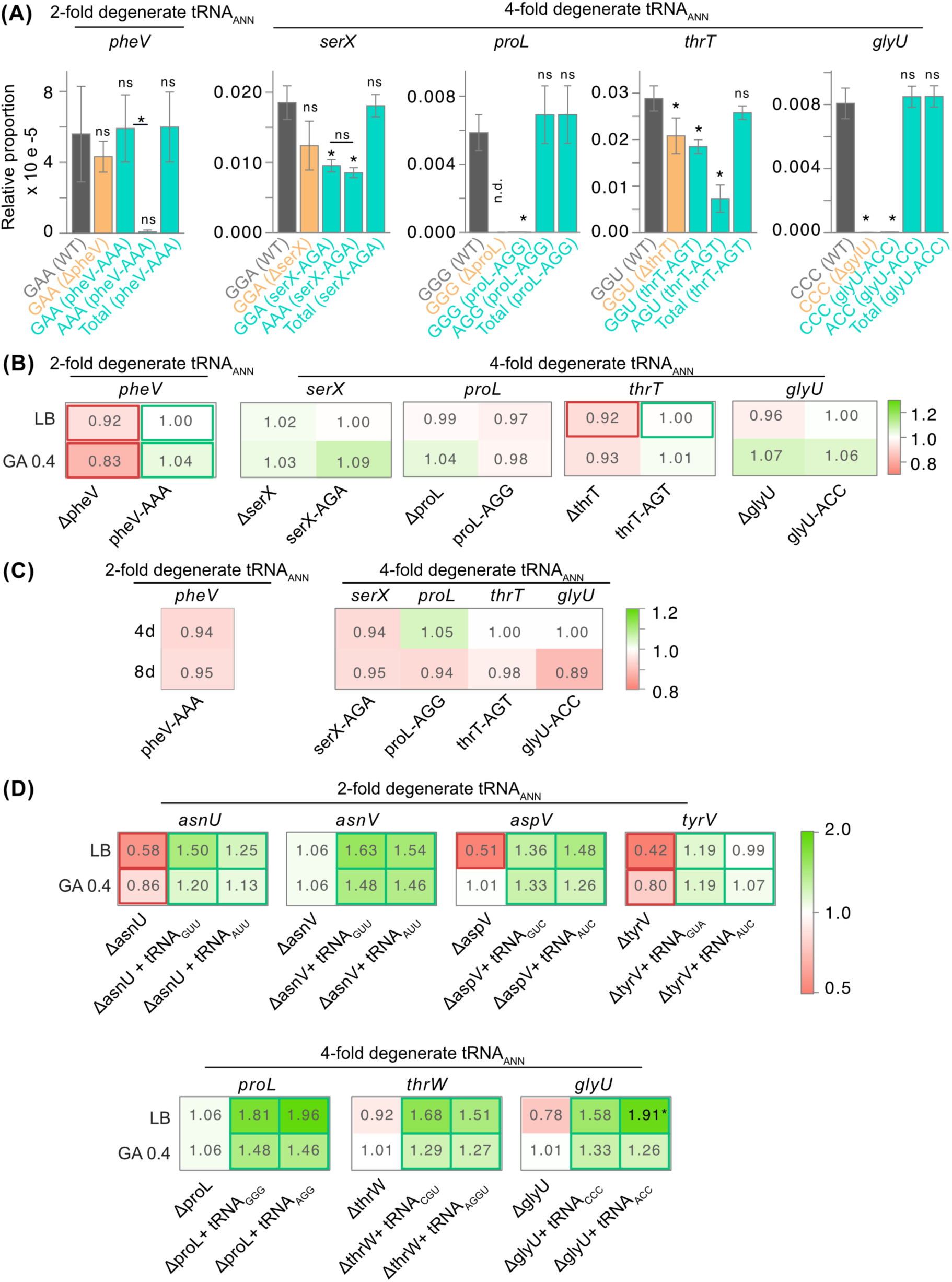
tRNA_ANN_ undergo normal maturation and can compensate for the loss of native tRNAs. **(A)** For a subset of tRNA genes (*pheV*, *serX*, *thrT*, *glyU*), we quantified the relative proportion of each tRNA species within the mature tRNA pool, for strains lacking the tRNA_BNN_ gene (KO, in orange), strains carrying a B34A substitution in the same tRNA_BNN_ gene (ANN strains, in green), and the wild type (WT, in grey). For *proL,* we quantified the relative proportion of each tRNA species for proL-ANN and WT. Bar graphs show the relative abundance of tRNA_BNN_ and tRNA_ANN_ of interest (n=3, mean ± standard deviation). Labels on x-axis indicate the anticodon detected, with the strain in parentheses. ‘Total’ indicates the combined abundance of tRNA_BNN_ and tRNA_ANN_ in ANN strains. ‘n.d.’ indicates not determined (experimental data not collected). Outcomes of statistical comparisons against tRNA_BNN_ in WT (first grey bar) are shown above each bar (pairwise Welch’s t-tests; asterisks indicate p < 0.05 and ns indicates p ≥ 0.05). Additional relevant comparisons not involving WT are indicated by connecting lines. See the source data file for Fig. 2A for all statistical comparisons and Fig. S3 for proportions of all tRNA species. **(B)** Growth rates relative to WT, for tRNA KO strains and ANN strains with tRNA_ANN_ expression from the genome (shown in Fig. 2A). KO strains that are significantly different from WT and ANN strains significantly different from KO are indicated by thick borders (Mann-Whitney U test, p < 0.05). Green and red borders indicate significant increase and decrease in growth rate respectively. **(C)** Growth rates of the five ANN strains in LB after serial passaging by transferring 1% v/v every 24 hours for 8 days. Each cell shows the mean growth rate of six such populations (relative to WT ancestor) for each strain on the fourth and eighth day (4d and 8d). None of these growth rates were significantly different from WT (paired t tests, p > 0.05). **(D)** Growth rates relative to WT+pACDH (empty vector control), for KO strains (+pACDH), and KO strains complemented with pACDH-tRNA_BNN_ and pACDH-tRNA_ANN_ for a subset of genes (*asnU*, *asnV*, *aspV*, *tyrV*, *proL*, *thrW*, *glyU*). The KO strains that were significantly different from WT, and the complementation strains significantly different from the KO+pACDH, are indicated by thick borders (Mann-Whitney U test, p < 0.05). Asterisks on cells for KO strains complemented with tRNA_ANN_ indicate a significant difference from strains complemented with tRNA_BNN_ (Mann-Whitney U test, p < 0.05).

We reasoned that if mature tRNA_ANN_ are translationally active, the fitness of strains carrying the B34A substitution should be similar to WT, and tRNA_ANN_ should rescue any deleterious effect of losing the respective tRNA_BNN_. Indeed, whereas ΔpheV and ΔthrT showed lower growth rate than WT, the growth rates of pheV-AAA and thrT-AGT were comparable to WT (Fig. 2B). Strains ΔpheV and ΔthrT also showed a longer lag phase in LB, which was shortened in ANN strains (Fig. S4A). The final OD was largely indistinguishable (± 5%) from WT for most KO and ANN strains, and was rescued in thrT-AGT (Fig. S4B). Improved growth parameters of these ANN strains as compared to the respective KO strains suggested that tRNA_ANN_ functionally replaced the native tRNA_BNN_. In contrast, deleting *serX*, *proL* and *glyU* did not impair growth (Fig. 2B), so rescue due to tRNA_ANN_ was not relevant. However, tRNA_ANN_ expression could still be toxic; but we did not see evidence of this. Growth rates (Fig. 2B) and final ODs (Fig. S4A) of serX-AGA, proL-AGG, and glyU-ACC strains were indistinguishable from WT, suggesting that these tRNA_ANN_ were also well tolerated during exponential growth. Lag phase length (Fig. 4B) was also largely indistinguishable (except serX-AGA and glyU-ACC in GA, where strains lacking these genes and those with B34A mutation, both had a longer lag phase than WT). Finally, ANN strains passaged in LB for eight days (about 80 generations) also did not show any fitness costs, suggesting a lack of long-term fitness effects of B34A substitutions on the genome (Fig. 2C). These observations further suggested that tRNA_ANN_ can successfully replace tRNA_BNN_ on the genome without significant fitness costs.

To further validate that tRNA_ANN_ can compensate for the loss of the respective tRNA_BNN_, we introduced plasmid-borne tRNA_ANN_ and tRNA_BNN_ into four of the 2D codon and three of the 4D codon tRNA KO strains and analysed their growth in several media, where we had previously observed differential effects of tRNA KOs (Raval et al. 2023). Across all five growth media and seven tRNA_ANN_ tested (35 combinations), tRNA_ANN_ improved overall growth of the respective KOs (Fig. S5A). The lag phase was significantly shorter than the respective KOs for 25 of 35 combinations (Fig. S5B). All strains showed exponential growth in two nutrient-rich media (LB and M9GA0.4), where complementation by tRNA_ANN_ increased growth rate for all 14 combinations tested (Fig. 2D) and increased final OD for 10 of 14 combinations (Fig. S5C). Complementation by tRNA_ANN_ vs. the respective tRNA_BNN_ increased the fitness of KOs to a similar extent, further corroborating the functionality of tRNA_ANN_.

Thus, all five tRNA_ANN_ introduced on the genome were folded, matured, and could functionally replace their respective tRNA_BNN_; and seven tRNA_ANN_ expressed from a plasmid could also functionally replace their native tRNA_BNN_ genes. A priori, a B34A substitution is unlikely to render a tRNA translationally inactive as described earlier, and prior studies also suggest that such tRNA_ANN_ have translational activity (Borén et al. 1993; Biddle et al. 2016; Schmitt et al. 2018; Schmitt et al. 2024). Taken together, we concluded that tRNA_ANN_ are generally translationally active and do not show evidence of substantial toxicity.

### Overexpression of 2D tRNA_ANN_ is often neutral or deleterious, while overexpression of 4D tRNA_ANN_ is often neutral or beneficial

The observation that expression of translationally active tRNA_ANN_ did not impair fitness of WT (Fig. 2B-C) was puzzling since their persistent rarity suggested otherwise. While tRNA_ANN_ expression from a single copy on the genome closely mimics potential B34A substitutions in natural bacterial populations, the tRNA_ANN_ thus expressed may also get outcompeted by native tRNA_BNN_ produced by other gene copies, and may not contribute substantially to translation. To reconcile a lack of clear fitness effects (even after ca. 80 generations) and the rarity of tRNA_ANN_ in nature, we reasoned that the fitness effects of tRNA_ANN_ may be attenuated when they are expressed from a single copy but may accumulate over evolutionary time scales. If so, tRNA_ANN_ overexpressed from a multi-copy plasmid may accentuate fitness effects and allow them to be detected within the timeframe of our experiments. In preliminary experiments, we observed differential effects of a subset of tRNA_ANN_ overexpressed from a low-copy-number IPTG-inducible plasmid under permissive growth conditions (37°C, LB medium). Both the tested 2D codons, Asn-tRNA_AUU_ and Asp-tRNA_AUC_, reduced relative growth rate (R_rel_) across all four IPTG concentrations (with no significant correlation between IPTG concentration and R_rel_), and Asn-tRNA_AUU_ also lowered the final OD_600_ (Fig. S6). However, the three 4D codon tRNA_ANN_ tested were largely neutral with respect to growth rate and final OD_600_ (Fig. S6). Thus, this preliminary experiment suggested that the fitness impacts of tRNA_ANN_ — especially those belonging to 2D codon boxes — may be exaggerated under high expression.

To systematically test the effects of overexpression, we analysed the fitness effects of tRNAs carrying all theoretically possible ANN anticodons (Fig. S1) introduced into the WT background from a high copy plasmid. In line with a previous study showing media-dependent costs and benefits of tRNA genes (Raval et al. 2023), we assessed fitness across media with varying nutrient contents and carbon sources. In half the tRNA_ANN_-media combinations tested, there was no significant fitness effect (Fig. 3A-B, Fig. S7A, Fig. S8-9) and the most consistent effects were observed during early growth (Fig. 3A; lag phase). Three out of the eight tested 2D codon tRNA_ANN_ were deleterious: Phe-tRNA_AAA_, Cys-tRNA_ACA_ and His-tRNA_AUG_ significantly prolonged the lag phase (Fig. 3A) and reduced growth rate (Fig. S8B, S9A, Fig. S9C) in at least two of six media. Phe-tRNA_AAA_ and Cys-tRNA_ACA_ also reduced final OD in four of the six media (Fig. S7A, Fig. S8C, Fig. S9B). Tyr-tRNA_AUA_, Asn-tRNA_AUU_ and Asp-tRNA_AUC_ prolonged the lag phase (Fig. 3A, Fig. S8A) and reduced growth rate in at least one medium (Fig. S8B, Fig. S9A). In contrast, Ser-tRNA_ACU_ improved all three growth parameters across at least two media (Fig. 3A, Fig. S7A, Fig. S8, S9) and Ile-tRNA_AAU_ increased growth rate (Fig. S8B, Fig. S9A) and the final OD (Fig. S8C, S9B) in at least two media. The magnitude of fitness effects (Fig. 3C) was skewed, with an overall significant increase in the lag phase length and also lower final OD; whereas effects on growth rate were less asymmetric and small. Overall, these results suggested that the deleterious effects of 2D tRNA_ANN_ were larger and more consistent than beneficial effects, particularly during early growth; and on average, 2D tRNA_ANN_ are likely to be neutral or deleterious.

**Figure 3:**
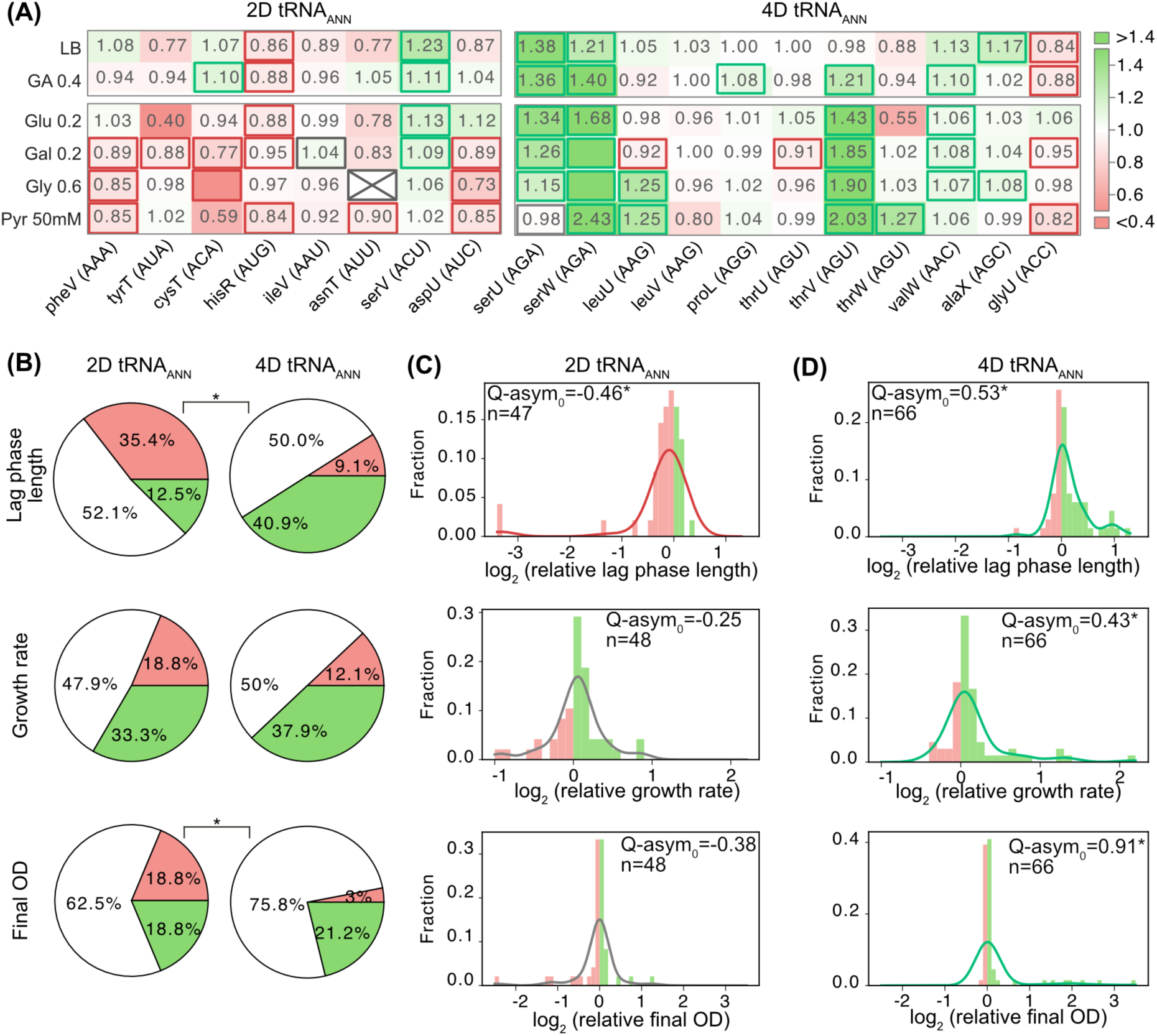
Fitness effects of overexpressing tRNA_ANN_ from a high-copy plasmid. **(A)** Heat maps show lag phase length of WT strains carrying tRNA_ANN_, relative to those carrying the corresponding native tRNA_BNN_ gene on the high copy plasmid pUC19 induced with 0.5mM IPTG at 37°C. Throughout, we compared fitness effects of overexpression of tRNA_ANN_ with that of respective tRNA_BNN_, calculating relative fitness as the growth parameter value of the strain expressing tRNA_ANN_ by that of the strain expressing tRNA_BNN_. Positive effects (tRNA_ANN_/ tRNA_BNN_ > 1) are shown in green and negative impacts (tRNA_ANN_/ tRNA_BNN_ < 1) in red. Significant differences between strains expressing tRNA_ANN_ and the corresponding tRNA_BNN_ (Mann-Whitney U test, p < 0.05; N = 4) with effect size ≥5% are highlighted by a thick border. Statistically significant differences with effect size < 5% (potentially reflecting noise from detection limits and fitting of the growth equation) are indicated by thick grey borders. The first two rows include nutrient-rich media and the next four rows include nutrient-poor media. In some cases, data for one or more replicates from either the tRNA_ANN_ strain or the respective tRNA_BNN_ variant did not fit the exponential growth equation whereas the other variant grew; e.g., Cys-tRNA_ACA_ in Gly 0.6 medium did not grow, unlike Cys-tRNA_GCA_ (raw OD_600_ vs. time curves are shown in Fig. S7). Such cases yielded infinite or infinitesimal values for growth parameters, so the corresponding cells in the heatmaps are empty; but the fill colours and outlines indicate the qualitative direction of relative growth (green if only tRNA_ANN_ strain grew and red if only tRNA_BNN_ strain grew). A cell is crossed out if neither the tRNA_ANN_ nor tRNA_BNN_ strain grew sufficiently well to calculate lag phase (e.g., tRNAs encoded by *asnT* in Gly 0.6). The absolute values of growth parameters are shown in Fig. S8 and summarised in the source data file for Fig. S8. **(B)** Fraction of cases where 2D and 4D tRNA_ANN_ show positive (green), negative (red) or neutral (white) effects for each growth parameter, compiling data across media from the heatmaps in Fig. 3A and Fig. S9A-B. Fisher’s exact tests were performed for each growth parameter to test whether 2D codon tRNA_ANN_ are more likely to be deleterious than 4D codon tRNA_ANN_. Asterisks indicate significant differences (p < 0.05). **(C-D)** Distribution of the mean relative fitness effect shown in the heatmaps in Fig. 3A and Fig. S9A-B, after log_2_ transformation (bandwidth= 0.1). Smoothed lines (kernel density bin width = 0.4) indicate the estimated underlying probability distribution. Q-asym_0_ estimates the magnitude of fitness effect skew towards beneficial (+1) to harmful (−1). Asterisks indicate a statistically significant skew, red/green denote harmful/beneficial skew, and grey denotes no significant skew (see Methods for calculation of Q-asym_0_ and statistical analysis).

We next investigated fitness effects of overexpressing 4D codon tRNA_ANN_, which did not show deleterious effects in our preliminary experiment. Six of the eleven 4D codon tRNA_ANN_ significantly improved lag phase length in at least two media, and four were neutral (Fig. 3A, Fig. S7B, Fig. S8A), in contrast to the results for 2D tRNA^ANN^ (Fig. 3A, Fig. S7A, Fig. S8A). Exponential growth rate was also improved by five 4D tRNA_ANN_ and remained unaffected by overexpression of four other 4D codon tRNA^ANN^ (Fig. S8B, S9A). Three of the 4D codon tRNA_ANN_ improved final OD and the remaining eight appeared to be neutral (Fig. S8C, Fig. S9B). Val-tRNA_AAC_ reduced growth rate in three media (Fig. S8B, Fig. S9A) and Gly-tRNA_ACC_ (the only universally absent 4D codon tRNA_ANN_) impaired lag phase as well as growth rate (Fig. 3A, Fig. S8A-B, Fig. S9A-B). Overall, apart from the universally absent Gly-tRNA_ACC_, 4D codon tRNA_ANN_ either did not affect growth or improved it (Fig. 3B, Fig. S9C). Both 4D Ser-tRNAs improved early and exponential growth in at least half of the media tested, whereas only one of the two Thr-tRNA_ANN_ was beneficial, suggesting backbone specific differences. Similar to 2D tRNA_ANN_, in half of the media-tRNA_ANN_ combinations tested, there were no significant fitness effects. In the remaining half of the cases tested, 4D codon tRNA_ANN_ also showed skewed fitness magnitudes, but opposite to that of 2D tRNA_ANN_. Overexpression of 4D tRNA_ANN_ skewed each of the three growth parameters significantly towards more beneficial effects (Fig. 3D). Thus, we concluded that on average, 4D tRNA_ANN_ are likely to be neutral or substantially beneficial, in contrast to 2D codon tRNA_ANN_ which were more likely to impair early growth (lag phase) and less likely to improve later stages (exponential growth and final OD) (Fig. 3B, Fig. S9C).

The differential fitness effects of 2D and 4D tRNA_ANN_ prompted us to consider the degeneracy of the genetic code and a prior observation that tRNA_ANN_ can superwobble, i.e., read all four codons in a box, in bacteria, mitochondria and eukaryotic cytosol (Sibler et al. 1986; Andachi et al. 1987; Borén et al. 1993; Inagaki et al. 1995; Watanabe et al. 1997; Von Nickisch-Rosenegk et al. 2001; Chen et al. 2002; Aldinger et al. 2012; Yokobori et al. 2013; Soma et al. 2023; Kompatscher et al. 2024; Schmitt et al. 2024). We speculated that a 2D tRNA_ANN_ could result in mistranslation because the tRNA will still be charged with the cognate amino acid but it can decode the non-cognate codons via superwobble (Fig. S10A). In contrast, a 4D codon tRNA_ANN_ will incorporate the same amino acid even if it decodes all four codons in the same box (Fig. S10B). Although direct measurement of mistranslation by 2D codon tRNA_ANN_ will require more detailed studies, we indirectly tested for this effect by using genome-wide codon usage to estimate the genome-wide mistranslation likelihood for each 2D codon tRNA_ANN_ (Fig. S10C). We expected that higher mistranslation likelihood should correspond to lower fitness in our experiments. Indeed, 2D codon Phe-tRNA_AAA_, Cys-tRNA_ACA_, Asn-tRNA_ATT_ with high mistranslation likelihood impaired fitness whereas Ser-tRNA_ACU_ and Ile-tRNA_AAU_ with low mistranslation likelihood improved it. This yielded a weak negative correlation between the overall fitness effect of each tRNA_ANN_ and its mistranslation likelihood (Fig. S10D), though we acknowledge that this analysis is constrained because only eight tRNA_ANN_ carrying 2D anticodon are theoretically possible. Nevertheless, this at least qualitatively suggested that mistranslation resulting from superwobbling may contribute to the negative fitness effects of 2D codon tRNA_ANN_. Superwobbling by 4D tRNA_ANN_ on the other hand, may be tolerated or even exploited due to the degeneracy of the genetic code.

### B34A substitutions are generally rare, but are better tolerated in 4D codon box tRNAs

The beneficial and neutral fitness effects of 4D codon tRNA_ANN_ were unexpected due to their near-universal absence, with the exception of two 4D codon tRNA_ANN_ in *Leuconostocaceae* (*Leuconostoc* and *Enococcus* sp.) reported in previous analyses of a relatively small number of bacterial genomes (Diwan and Agashe 2018; Ehrlich et al. 2021). We re-evaluated the rarity of tRNA_ANN_ using the entire prokaryotic tRNA repertoire known to date, which includes 246,393 predicted tRNA sequences across 4047 bacteria and 10,517 tRNAs from 220 archaea in the GtRNA database (Chan and Lowe 2016; Thornlow et al. 2020). Across 10,517 predicted archaeal tRNAs, only three cases of tRNA_ANN_ were found, all from 4D codon boxes (Fig. 4A). Of 8383 bacterial tRNA_ANN_ genes, 7,688 encoded Arg-tRNA_ACG_, supporting the rarity of other ANN anticodons across bacteria (Fig. 4A). However, the 695 non-ACG tRNA_ANN_ sequences did include all other theoretically possible ANN. Notably, 81% of these were 4D codon tRNA_ANN_, with Leu-tRNA_AAG_ and Thr-tRNA_AGT_ being most frequent (Fig. 4B). Of the remaining (19%) 2D tRNA_ANN_ sequences, half were Phe-tRNA_AAA_ and His-tRNA_AUG_, which are potentially compatible with the A-to-I modification system.

**Figure 4:**
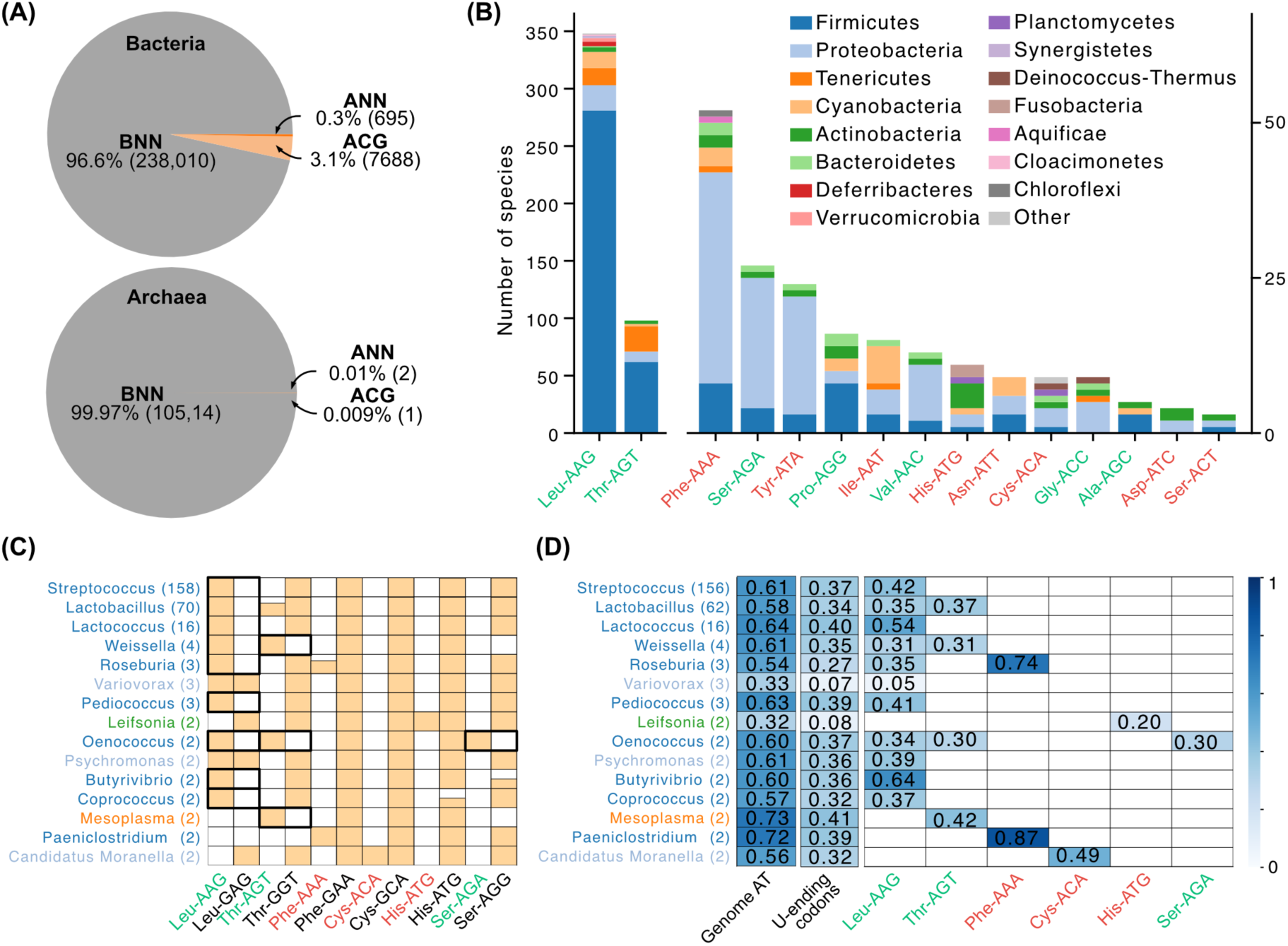
Occurrence of tRNA_ANN_ across prokaryotes. **(A)** Fractions of tRNA_ANN_ and tRNA_BNN_ within all bacterial and archaeal tRNAs reported in the GtRNA DB. The number of tRNA genes is indicated in parentheses. **(B)** Number of bacterial genomes carrying each tRNA_ANN_ gene, grouped by phylum. 4D and 2D tRNA_ANN_ are labelled in green and red, respectively. For tRNA_ANN_ present in fewer than 50 species, refer to the right-hand Y axis. **(C)** Bacterial genera (colored by phylum, see panel B) where at least one tRNA_ANN_ is found in all species (number of species is indicated in parentheses). The coloured portion of each cell represents the fraction of species where the tRNA_ANN_ is present. 4D and 2D tRNA_ANN_ are labelled in green and red, respectively. For comparison, presence of the respective tRNA_GNN_ is also shown (labelled in black). Instances where a tRNA_NN_ potentially replaced a tRNA_GNN_ (indicated by a concomitant lack of the respective tRNA_GNN_) are highlighted by black squares. **(D)** For the genera shown in panel C, average genome AT content (1 indicating 100% AT), fraction of U-ending codons out of all codons used in the genome, and fraction of U-ending codons within the codon box where the tRNA_ANN_ is present (averaged across all species within a genus).

Firmicutes encoded the highest number of tRNA_ANN_ (Fig. 4B, Fig. S10), and within this phylum, all *Lactobacillus*, *Lactococcus* and *Streptococcus* species encoded Leu-tRNA_AAG_ (Fig. 4C). These three genera lacked Leu-tRNA_GAG_, suggesting a G34A substitution dating back at least until their common ancestor (Fig. 4C). Thr-tRNA_AGT_ also showed a similar pattern, albeit for fewer genera. The presence of an ANN anticodon in lieu of a GNN is similar to Arg-tRNAs where ACG is preferred and GCG is absent, and suggests that these genera not only encode, but potentially prefer these specific 4D tRNA_ANN_. Interestingly, these two ANNs only replaced GNNs, but not other BNN isoacceptors, suggesting that potential superwobbling by unmodified ANNs may be suboptimal even in 4D codon boxes (e.g., due to slower translation rate). In contrast, 2D-codon tRNA_ANN_ were present sporadically across species from different genera (Fig. S11) and always in the presence of tRNA_GNN_ (Fig. S12). We also found some tRNA_ANN_ genes in Proteobacteria (genera *Salmonella*, *Escherichia*) and Tenericutes (subphylum Mollicutes), although their occurrence was scattered within the subphyla or genera, suggesting lineage specific acquisition or loss (Fig. 4B, S11).

Next, we analysed genomic features of species that do have tRNA_ANN_. The 15 genera with at least one tRNA_ANN_ found in all species (i.e., candidates for acquisition by their respective ancestors) all had high genomic AT content ranging from 60-70% (Fig. 4D) and 30-40% of all codons in these genomes were U-ending. In genera such as *Streptococcus* and *Lactococcus*, U-ending codons were also preferred over their synonyms in the same codon box where tRNA_ANN_ was present. However, in other genera, this preference was only moderate, suggesting that factors other than codon usage contribute to the retention of tRNA_ANN_ in some lineages. Lastly, 88% of the species with at least one non-ACG tRNA_ANN_ encode a TadA homologue (Fig. S11) which may modify these tRNA_ANN_ (e.g., Leu-tRNA_AAG_ is modified in some bacteria (Rafels-Ybern et al. 2019; Wulff et al. 2024; Katanski et al. 2026).

Such large-scale genome-based predictions are somewhat limited due to prediction errors and a lack of direct evidence for gene expression and functionality. We therefore retested the broad patterns after applying more stringent filters for tRNA gene prediction using gene score ≥70 bits (a measure of the similarity of the gene sequence to other tRNA genes, and the likelihood of the product folding like a tRNA) on gtRNA-db (last accessed on 21.08.2026, via the tool tRNA-exporter; see Table S2). We now detected only 132 tRNA_ANN_ genes (instead of 695 pre-filter, out of ca. 240K bacterial tRNA genes). Among these, 4D tRNA_ANN_ genes were still more abundant than 2D tRNA_ANN_ and were limited to specific taxa (127 out of 132 are 4D, 90 of which are Thr-tRNA_AGT_ and 28 are Leu-tRNA_AAG_ predicted in *Lactobacillus*, *Leuconostoc* and *Mycoplasma*). Thus, non-ACG tRNA_ANN_ are indeed rare, and among the exceptions, 4D codon tRNA_ANN_ (e.g., Leu-tRNA_GAG_ and Thr-tRNA_AGT_) are retained across evolutionary timescales and potentially even preferred in some genera. Overall, other 4D codon tRNA_ANN_ are generally better tolerated than unmodified 2D codon tRNA_ANN_, corroborating our experimental results.

## DISCUSSION

The near-universal absence of unmodified tRNAs with A34 (tRNA_ANN_) decoding both two-fold (2D) and four-fold (4D) degenerate codon boxes imply strong purifying selection, arguably acting since early cellular evolution. Although numerous sources of purifying selection can be speculated (Fig. 1B), a key first step towards explaining the absence of tRNA_ANN_ is to directly determine the functionality and fitness effects of expressing such tRNAs. Our results show that tRNA_ANN_ in their native backbones are tolerated, folded, matured and translationally active in *E.* coli. A majority of the 4D codon tRNA_ANN_ were also tolerated in an unmodified state and showed neutral or positive fitness effects when overexpressed. However, 2D codon box tRNA_ANN_ were more likely to be deleterious when overexpressed and their magnitude of fitness effect appeared to scale with a coarsely estimated likelihood of mistranslation. Furthermore, all tested 4D codon tRNA_ANN_ were tolerated unmodified, whereas the only tested 2D codon Phe-tRNA_AAA_ was modified to Phe-tRNA_IAA_. The INN anticodon can decode only three codons (Oba et al. 1991; Curran 1995; Gerber and Keller 1999; Wolf et al. 2002; Yokobori et al. 2013; Schmitt et al. 2024) and thus it is likely to mis-incorporate amino acids at fewer codons than unmodified ANN, which can superwobble, i.e. read all four codons within a codon box (Sibler et al. 1986; Andachi et al. 1987; Borén et al. 1993; Inagaki et al. 1995; Watanabe et al. 1997; Von Nickisch-Rosenegk et al. 2001; Chen et al. 2002; Aldinger et al. 2012; Yokobori et al. 2013; Soma et al. 2023; Kompatscher et al. 2024; Schmitt et al. 2024). A prior study also found that a 2D codon His-tRNA_AUG_ underwent such modification (Biddle et al. 2016; Schmitt et al. 2024). The enzyme responsible for I34 modification of these “novel” bacterial tRNA_ANN_ substrates remains unknown, though a parsimonious explanation is TadA, which canonically modifies Arg-tRNA_ACG_ and was recently shown to modify numerous mRNAs in *E. coli* (Arad et al. 2026). Our study thus suggests potential substrate flexibility of this ancient and essential enzyme (Wolf et al. 2002; Delannoy et al. 2009; Diwan and Agashe 2018; Torres et al. 2021) that may mitigate the deleterious effects of other tRNA_ANN_ and confer additional advantages, though this remains to be confirmed. Intriguingly, the two tRNA_ANN_ that are compatible with modification also constituted 50% of the rarely found 2D codon tRNA_ANN_ across bacteria (Fig. 4B), suggesting that A-to-I modification might mitigate mistranslation and fitness impairment by such tRNAs. Taken together, selection for translational fidelity hence emerges as one plausible explanation for the absence of 2D codon tRNA_ANN_. Fitness defects could arise due to direct misincorporation at 2D codon tRNA_ANN_ or an increase in other forms of mistranslation such as frameshifting, or indirectly due to disruptions in inosine modification; much more work is needed to distinguish between these and other mechanisms.

In contrast to 2D codon tRNA_ANN_, superwobbling by a tRNA_ANN_ within a 4D codon box is not expected to cause mistranslation. Indeed, we found that all 4D tRNA_ANN_ (except Gly-tRNA_ACC_) were neutral or substantially beneficial. 4D tRNA_ANN_ were also more likely to be retained or even preferred as compared to 2D codon tRNA_ANN_ among the exceptional cases of A34 occurrences across bacteria. Nevertheless, seven of eight 4D tRNA_ANN_ are missing from 99.99% of predicted bacterial tRNA repertoires. Both bacteria and eukaryotes decode the arginine 4D codon box with Arg-tRNA_ACG_, whereas for the other seven 4D codon boxes, bacteria prefer tRNA_UNN_ and eukaryotes use tRNA_ANN_ (Novoa et al. 2012). Each of these 4D codon tRNAs is modified via A34-to-I34 or U34-to-cmo^5^U34 (Novoa et al. 2012; Diwan and Agashe 2018) and tRNA_GNN_ alternatives are absent. Codons recognized by these modified tRNAs are enriched in highly expressed genes, and these tRNAs (Novoa et al. 2012) and A-to-I modification appear to be essential (Wolf et al. 2002; Delannoy et al. 2009; Torres et al. 2021), further underscoring a crucial role of tRNA modification in decoding 4D codon boxes. Indeed, it has been proposed that coevolution between MEs and tRNA repertoires shaped the ancient preference for modifiable 4D codon tRNA_UNN_ in bacteria and tRNA_ANN_ in eukaryotes (Novoa et al. 2012). Unmodified 4D codon tRNA_GNN_ (absent across eukaryotes and present with G34-to-queuosine modification in most bacteria (Diwan and Agashe 2018)) makes the second and third bases error-prone, causing mistranslation outside the cognate codon boxes, and resulting in cellular toxicity (Pernod et al. 2021). Another recent study shows that A34 modification interacts with neighboring bases to influence translation fidelity under stress (Raval et al. 2026). These studies underscore the idea that specific anticodons — potentially also ANN — in specific backbones are inherently error-prone and thus, selection for translational fidelity may also contribute to the absence of 4D tRNA_ANN_. However, the neutral and positive fitness effects of unmodified 4D tRNA_ANN_ observed here require future studies to mechanistically reconcile them with their absence in nature.

While mistranslation is generally thought to be deleterious (O’connor et al. 1992; Beebe et al. 2003; Bacher et al. 2004; Kohanski et al. 2008; Berg et al. 2019; Kelly et al. 2019), it can also be beneficial under specific conditions, and especially under stress (Fan et al. 2015; Samhita et al. 2020; Samhita et al. 2021; Samhita 2022). The extent of mistranslation also varies across tRNA_ANN_ due to differences in stability across superwobble base-pairs (e.g., A-A is least stable). Consequences of amino acid misincorporation on protein structure and fitness also depends on the physico-chemical differences between the exchanged amino acids; thus, misincorporation by some tRNA_ANN_ (e.g., replacing Lys with Asn) may be less detrimental than others (e.g., Arg to Ser) (Sengupta et al. 2007). Fitness effects of mistranslation are therefore likely to vary across specific tRNA_ANN_ and environments, and our suggestion that mistranslation contributes to the absence of tRNA_ANN_ remains to be directly tested. A first step would be to measure the extent of mistranslation in cells expressing tRNA_ANN_, e.g., via mass-spectrometric analysis of a reporter protein. Strains with high-fidelity ribosomes (Ruusala et al. 1984; Chumpolkulwong et al. 2004; Agarwal et al. 2015) and downregulation of *tadA* could also be used to test mistranslation-driven fitness defects, which should be alleviated and exacerbated in the respective strain backgrounds. Likewise, antibiotics reducing ribosomal fidelity should also amplify such fitness defects. Lastly, we note that despite our normalization of fitness effects of overexpressed tRNA_ANN_ with that of overexpressed tRNA_BNN_, we cannot rule out that overexpression of tRNA_ANN_ caused extremely high mistranslation or resulted in other effects qualitatively different from those observed in nature. Thus, such overexpression may not have amplified natural fitness effects as we had intended. We hope that future studies using more sensitive fitness assays and long-term evolution may provide further insights.

Overall, while mistranslation is an attractive hypothesis, given these causes of substantial variation in its effects across specific tRNA_ANN_ and environments, it is unlikely to be the sole explanation for the rarity of all tRNA_ANN_ across prokaryotes. Therefore, we speculate important contributions from other mechanisms, likely as universal as core features of the translation machinery. For instance, correct pairing at the A site in bacterial ribosomes causes A1492/A1493 to flip out from 16S rRNA into the minor groove of the first two codon-anticodon base pair and promotes EF-Tu GTP hydrolysis and amino acylated tRNA accommodation (Schmeing and Ramakrishnan 2009). It is possible that an adenosine at the first anticodon position causes steric hindrance (due to the presence of two additional As in the vicinity) and impairs this essential step inside the decoding center, which is conserved across the tree of life (Rodnina and Wintermeyer 2009; Schmeing and Ramakrishnan 2009; Dever et al. 2018; Tirumalai et al. 2021). However, an adenosine is tolerated on the mRNA (e.g., in A-ending codons), suggesting that any steric hindrance is likely specific to the accommodation tRNA with A34. While this hypothesis remains to be tested via structural or molecular dynamics analyses, it serves as a useful example of the nature of potential fundamental explanations for the ancient and persistent rarity of tRNA_ANN_.

Growth phase, growth medium and anticodon specific fitness effects of distinct tRNA_ANN_ might also suggest additional factors contributing to their absence. For instance, tRNA_ANN_ had stronger fitness effects in amino acid-poor media and during early growth, whereas most, particularly 4D tRNA_ANN_, were neutral in nutrient-rich media and in the stationary phase. This suggests stronger selection on tRNA_ANN_ under resource limitation and/or high translational demand. Such conditions could help eliminate deleterious tRNA_ANN_ (e.g. some 2D tRNA_ANN_), but should favour beneficial ones (e.g. most 4D tRNA_ANN_), deepening the puzzle of their absence and pointing to additional constraints not captured by these assays. Fitness assays under nutrient fluctuation, environmental stress, or competition with WT may reveal further costs of unmodified 4D tRNA_ANN_. Another interesting pattern in our data is that among the 2D and 4D tRNA_ANN_, anticodons in which A34 was followed by A35 or U35, resulting in an A–U/U–A pair at the second codon position, were more detrimental than those containing C35 or G35. This trend was particularly apparent among the 2D tRNA_AAA_, tRNA_AUA_, tRNA_AUG_ that tended to impair early growth, while tRNA_AAU_, tRNA_AUU_, and tRNA_AUC_ also showed detrimental effects on lag phase, although these were not statistically significant. However, the C35-containing 2D tRNA_ACA_ was also detrimental, whereas tRNA_ACU_ was beneficial. Thus, base-pair strength at position 35 may contribute to the fitness effects of B34A substitutions; but likely involve additional factors.

In closing, the near-complete absence of unmodified tRNA_ANN_ remains intriguing, more so given the largely neutral and sometimes beneficial effects that we observed. At a population size of a billion cells and assuming a 10% fitness advantage (i.e., selection coefficient *s*=0.1), a mutation creating a tRNA_ANN_ from a tRNA_BNN_ has about 17% chance of reaching fixation in an average of 410 generations, as inferred from 1000 discrete-time Wright–Fisher transitions (Fig. S13). Even a tRNA_ANN_ with a 10% disadvantage can persist for tens of generations, and a tRNA_ANN_ with a disadvantage smaller than 1% can persist for over a thousand generations. Nonetheless, tRNA_ANN_ are remarkably rare. Although cases with negative fitness effects of 2D tRNA_ANN_ suggest that selection for translational fidelity likely contributes to their absence, the neutral and beneficial effects observed for 4D tRNA_ANN_ suggests that other fundamental constraints on these molecules, operating in nature, remain hidden.

## DATA AVAILABILITY

Supplementary figures are available with this submission. Raw reads for YAMAT-seq are available on from NCBI GEO; accession number GSE328815. Additional data are available on Zenedo (10.5281/zenodo.23323378) which includes, but is not limited to, the following supplementary data and in-house scripts:

Source data for the main and supplementary figures

Table S1 (Primers and strains from this study)

Table S2 (tRNA_ANN_ genes across prokaryotes after filtering for Gene Scores on GtRNAdb)

Scripts, Source files for the scripts, Output files and plots.

## ACKNOWLEDGEMENTS

We thank Saurabh Mahajan, Laasya Samhita, Supratim Sengupta and members of the Agashe lab for discussion and critical comments on the manuscript; the NCBS NGS facility for help with genome and YAMAT sequencing; Gaurav Diwan and Joshua Miranda for setting up and maintaining our automated growth measurement system; Gunda Dechow-Seligmann and Sven Künzel for helping with data collection for YAMAT-seq; George Stoletov, Raagini Biswas, Adrita Chakraborty and Pratibha Sanjenbam for helping with the experiments; and the NCBS laboratory kitchen staff for their crucial support. PKR acknowledges the use of ChatGPT (OpenAI) for assistance with writing Python scripts. We acknowledge funding and support from the National Centre for Biological Sciences (NCBS-TIFR) and the Department of Atomic Energy, Government of India (Project Identification No. RTI 4006) to DA, a CSIR-UGC-NET June/2018/430 fellowship to PKR, the Max Planck Society (JG and SL), and the International Max Planck Research School for Evolutionary Biology (SL).

## AUTHOR CONTRIBUTIONS

PKR: Conceptualization, Experimental design, Methodology, Investigation, Data curation, Validation, Formal analysis, Visualization, Writing - original draft, review and editing. SL: Investigation, Data curation, Formal analysis JG: Experimental design, Funding acquisition, Resources, Methodology, Writing - review and editing. DA: Conceptualization, Project administration, Supervision, Experimental design, Funding acquisition, Resources, Writing - review and editing.

