## Supplementary Figures 1-13 for "Rare tRNAs with A-starting anticodons are functional and show codon-box dependent fitness effects"

**Figure S1: Introduction of $tRNA_{ANN}$ into *E. coli.*** Genome wide codon usage (from https://www.kazusa.or.jp/codon/) and tRNA gene copy numbers (from http://gtrnadb.ucsc.edu/index.html) in *E. coli* K-12 MG1655, consolidated as per the key on the bottom right. tRNAs with Adenosine 34 ($tRNA_{ANN}$) are shown in red. Native tRNA genes that were mutated to $tRNA_{ANN}$ in this study are indicated in bold. ANNs expressed from a plasmid are underlined, and those expressed from the native genomic location are indicated by asterisks.

| Amino acid | Codon (Codons per thousand) | Anticodon (tRNA gene copies) |
|---|---|---|
| Phe | UUU (19.7) | AAA (0) |
| Phe | UUC (15) | **AAG* (2)** |
| Leu | UUA (15.2) | AAU (1) |
| Leu | UUG (11.9) | AAC (1) |
| Ser | UCU (8.5) | AGA (0) |
| Ser | UCC (8.5) | **AGG* (2)** |
| Ser | UCA (7.1) | **AGU (1)** |
| Ser | UCG (8.9) | **AGC (1)** |
| Tyr | UAU (16.3) | AUA (0) |
| Tyr | UAC (12.8) | **AUG (3)** |
| Stop | UAA (2.0) | AUU |
| Stop | UAG (0.2) | AUC |
| Cys | UGU (5.9) | ACA (0) |
| Cys | UGC (8) | **ACG (1)** |
| Stop | UGA (0.9) | ACU |
| Trp | UGG (15.3) | ACC (1) |
| Leu | CUU (11.9) | GAA (0) |
| Leu | CUC (10.5) | **GAG (1)** |
| Leu | CUA (5.3) | GAU (1) |
| Leu | CUG (46.9) | **GAC (4)** |
| Pro | CCU (7.0) | GGA (0) |
| Pro | CCC (5.5) | **GGG* (1)** |
| Pro | CCA (8.5) | GGU (1) |
| Pro | CCG (23.2) | GGC (1) |
| His | CAU (12.9) | GUA (0) |
| His | CAC (9.7) | **GUG (1)** |
| Gln | CAA (15.7) | GUU (2) |
| Gln | CAG (28.8) | GUC (2) |
| Arg | CGU (7.0) | GCA (4) |
| Arg | CGC (5.5) | GCG (0) |
| Arg | CGA (8.5) | GCU (0) |
| Arg | CGG (23.2) | GCC (1) |
| Ileu | AUU (30.5) | UAA (0) |
| Ileu | AUC (18.2) | **UAG (3)** |
| Ileu | AUA (3.7) | UAU (0) |
| f/Met | AUG (24.8) | UAC (4/2) |
| Thr | ACU (8.9) | UGA (0) |
| Thr | ACC (23.3) | **UGG* (2)** |
| Thr | ACA (7.0) | **UGU (1)** |
| Thr | ACG (14.37) | **UGC (2)** |
| Asn | AAU (17.6) | UAA (0) |
| Asn | AAC (21.6) | **UUG (4)** |
| Lys | AAA (33.6) | UUU (6) |
| Lys | AAG (10.2) | UUC (0) |
| Ser | AGU (8.7) | UCA (0) |
| Ser | AGC (16.0) | **UCG (1)** |
| Arg | AGA (2.0) | UCU (1) |
| Arg | AGG (1.2) | UCC (1) |
| Val | GUU (16.8) | CAA (0) |
| Val | GUC (11.7) | **CAG (2)** |
| Val | GUA (11.5) | CAU (5) |
| Val | GUG (26.4) | CAC (0) |
| Ala | GCU (15.3) | CGA (0) |
| Ala | GCC (25.5) | **CGG (2)** |
| Ala | GCA (20.2) | CGU (3) |
| Ala | GCG (33.6) | CGC (0) |
| Asp | GAU (32.2) | CUA (0) |
| Asp | GAC (19.0) | **CUG (3)** |
| Glu | GAA (39.5) | CUU (0) |
| Glu | GAG (17.8) | CUC (4) |
| Gly | GGU (24.8) | CCA (0) |
| Gly | GGC (29.4) | CCG (4) |
| Gly | GGA (7.9) | CCU (1) |
| Gly | GGG (10.9) | **CCC* (1)** |

Key:

| Amino acid | | |
|---|---|---|
| | Codon (5'-3') | (Codons per thousand) |
| | Anticodon (3'-5') | (tRNA gene copies) |

**Figure S2: Differences in the backbones of tRNAs decoding *E. coli* family codon boxes.** Alignments of tRNA isoacceptors (http://gtrnadb.ucsc.edu/index.html) with bases forming the D-arm, anticodon loop and T-arm highlighted in brick, green, light blue, respectively. Pairwise differences between each tRNA isoacceptor is shown above each alignment; tRNA$_{BNN}$ from which we derived tRNA$_{ANN}$ are in bold.

Serine (**Ser-tRNA$_{CGA}$** vs **Ser-tRNA$_{TGA}$** = 37%, Ser-tRNA$_{GGA}$ vs Ser-tRNA$_{CGA}$ = 34%, Ser-tRNA$_{GGA}$ vs Ser-tRNA$_{TGA}$ = 46%)

```
>>>>>>>..>>>............<<<.>>>>>.......<<<<<..>>>>>.........<<<<<..>>>>>.......<<<<<..<<<<<<<....
GGTGAGGTGTCCGAGTGGCtgA.AGGAGCACGCCTGGAAAGTGTGT.ATACG..GCAA...CGTAT.CGGGGGTTCGAATCCCCC..CCTCACCGCCA tRNA-Ser-GGA-1-1
GGTGAGGTGTCCGAGTGGCtgA.AGGAGCACGCCTGGAAAGTGTGT.ATACG..GCAA...CGTAT.CGGGGGTTCGAATCCCCC..CCTCACCGCCA tRNA-Ser-GGA-1-2
>>>>>>>..>>>............<<<.>>>>>.......<<<<<..>>>>>>.......<<<<<<..>>>>>.......<<<<<..<<<<<<<....
GGAGAGATGCCGGAGCGGCtgA.ACGGACCGGTCTCGAAAACCGGA.GTAGGG.GCAA..CTCTAC.CGGGGGTTCAAATCCCCC..TCTCTCCGCCA tRNA-Ser-CGA-1-1
>>>>>>>..>>>............<<<.>>>>>.......<<<<<..>>>>...........<<<<..>>>>>.......<<<<<..<<<<<<<....
GGAAGTGTGGCCGAGCGGTtgA.AGGCACCGGTCTTGAAAACCGGCGACCC...GAAA....GGGTTCCAGAGTTCGAATCTCTG..CGCTTCCGCCA tRNA-Ser-TGA-1-1
>>>>>>>..>>>............<<<.>>>>>.......<<<<<..>>>>>>>.....<<<<<<<..>>>>>.......<<<<<..<<<<<<<....
GGTGAGGTGGCCGAGAGGCtgA.AGGCGCTCCCCTGCTAAGGGAGT.ATGCGGTCAAAaGCTGCAT.CCGGGGTTCGAATCCCCG..CCTCACCGCCA tRNA-Ser-GCT-1-1
```

Leucine (**Leu-tRNA$_{GAG}$** vs **Leu-tRNA$_{CAG}$** = 33%, Leu-tRNA$_{GAG}$ vs Leu-tRNA$_{TAG}$ = 39%, Leu-tRNA$_{CAG}$ vs Leu-tRNA$_{TAG}$ = 36%)

```
>>>>>>>..>>>...........<<<.>>>>>.......<<<<<..>>>.>.........<.<<<...>>>>>.......<<<<<.<<<<<<<....
GCCGAGGTGGTGGAATTGGTaGACACGCTACCTTGAGGTGGTAGTGCCCAA.........TAGGGCtTACGGGTTCAAGTCCCGT.CCTCGGTACCA tRNA-Leu-GAG-1-1
>>>>.>>..>>>...........<<<.>>>>>.......<<<<<..>>>>...........<<<<...>>>>>.......<<<<<.<<.<<<<....
GCGAAGGTGGCGGAATTGGTaGACGCGCTAGCTTCAGGTGTTAGT.GTCC.TTAC......GGACG.TGGGGGTTCAAGTCCCCC.CCCTCGCACCA tRNA-Leu-CAG-1-1
GCGAAGGTGGCGGAATTGGTaGACGCGCTAGCTTCAGGTGTTAGT.GTCC.TTAC......GGACG.TGGGGGTTCAAGTCCCCC.CCCTCGCACCA tRNA-Leu-CAG-1-2
GCGAAGGTGGCGGAATTGGTaGACGCGCTAGCTTCAGGTGTTAGT.GTCC.TTAC......GGACG.TGGGGGTTCAAGTCCCCC.CCCTCGCACCA tRNA-Leu-CAG-1-3
GCGAAGGTGGCGGAATTGGTaGACGCGCTAGCTTCAGGTGTTAGT.GTTC.TTAC......GGACG.TGGGGGTTCAAGTCCCCC.CCCTCGCACCA tRNA-Leu-CAG-2-1
>>>>>>>..>>>...........<<<.>>>>>.......<<<<<..>>>.............<<<...>>>>>.......<<<<<.<<<<<<<....
GCGGGAGTGGCGAAATTGGTaGACGCACCAGATTTAGGTTCTGGC.GCC..GCAA.......GGTG.TGCGAGTTCAAGTCTCGC.CTCCCGCACCA tRNA-Leu-TAG-1-1
```

Threonine (**Thr-tRNA$_{GGT}$** vs **Thr-tRNA$_{CGT}$** = 31%, Thr-tRNA$_{GGT}$ vs **Thr-tRNA$_{TGT}$** = 35%, Thr-tRNA$_{CGT}$ vs Thr-tRNA$_{TGT}$ = 31%**)**

```
>>>>>>>..>>>>.........<<<<.>>>>>.......<<<<<................>>>>>.......<<<<<<<<<<<<....
GCTGATATAGCTCAGTTGGT.AGAGCGCACCCTTGGTAAGGGTGAG...........GtCGGCAGTTCGAATCTGCCTATCAGCACCA tRNA-Thr-GGT-1-1
GCTGATATGGCTCAGTTGGT.AGAGCGCACCCTTGGTAAGGGTGAG...........GtCCCCAGTTCGACTCTGGGTATCAGCACCA tRNA-Thr-GGT-2-1
>>>>>>>..>>>>.........<<<<..>>>>.......<<<<.................>>>>>.......<<<<<<<<<<<<....
GCCGATATAGCTCAGTTGGT.AGAGCAGCGCATTCGTAATGCGAAG...........GtCGTAGGTTCGACTCCTATTATCGGCACCA tRNA-Thr-CGT-1-1
>........>>>>.........<<<<.>>>>>.......<<<<<................>>>>>.......<<<<<......<....
G......TAGTTAAAAATGCaTTAACATCGCATTCGTAATGCGAAG...........GtCGTAGGTTCGACTCCTATTATCGGCACCA tRNA-Thr-CGT-2-1
>>>>>>>..>>>>.........<<<<.>>>>>.......<<<<<................>>>.>.......<.<<<<<<<<<<....
GCCGACTTAGCTCAGTAGGT.AGAGCAACTGACTTGTAATCAGTAG...........GtCACCAGTTCGATTCCGGTAGTCGGCACCA tRNA-Thr-TGT-1-1
```

**Figure S3. Fold change in the relative abundance of each tRNA species in experimental strains.** For a subset of strains, we quantified the relative proportion of each tRNA species within the mature tRNA pool. We tested strains lacking a $tRNA_{BNN}$ gene ('KO'; *pheV*, *serX*, *thrT*, or *glyU*) and corresponding strains carrying the G34A substitution in the same gene (i.e., $tRNA_{ANN}$). For *proL,* we were unable measure the KO strain, so data are shown only for proL-ANN. Each cell shows fold change of a tRNA species (each row) in a given strain (each column) relative to the wild-type control included in the same experiment (n = 3 biological replicates per strain). Cases where tRNA abundance differs significantly between the test strain and wild type are marked by a bold cell border (Welch's t-test, $p < 0.05$).

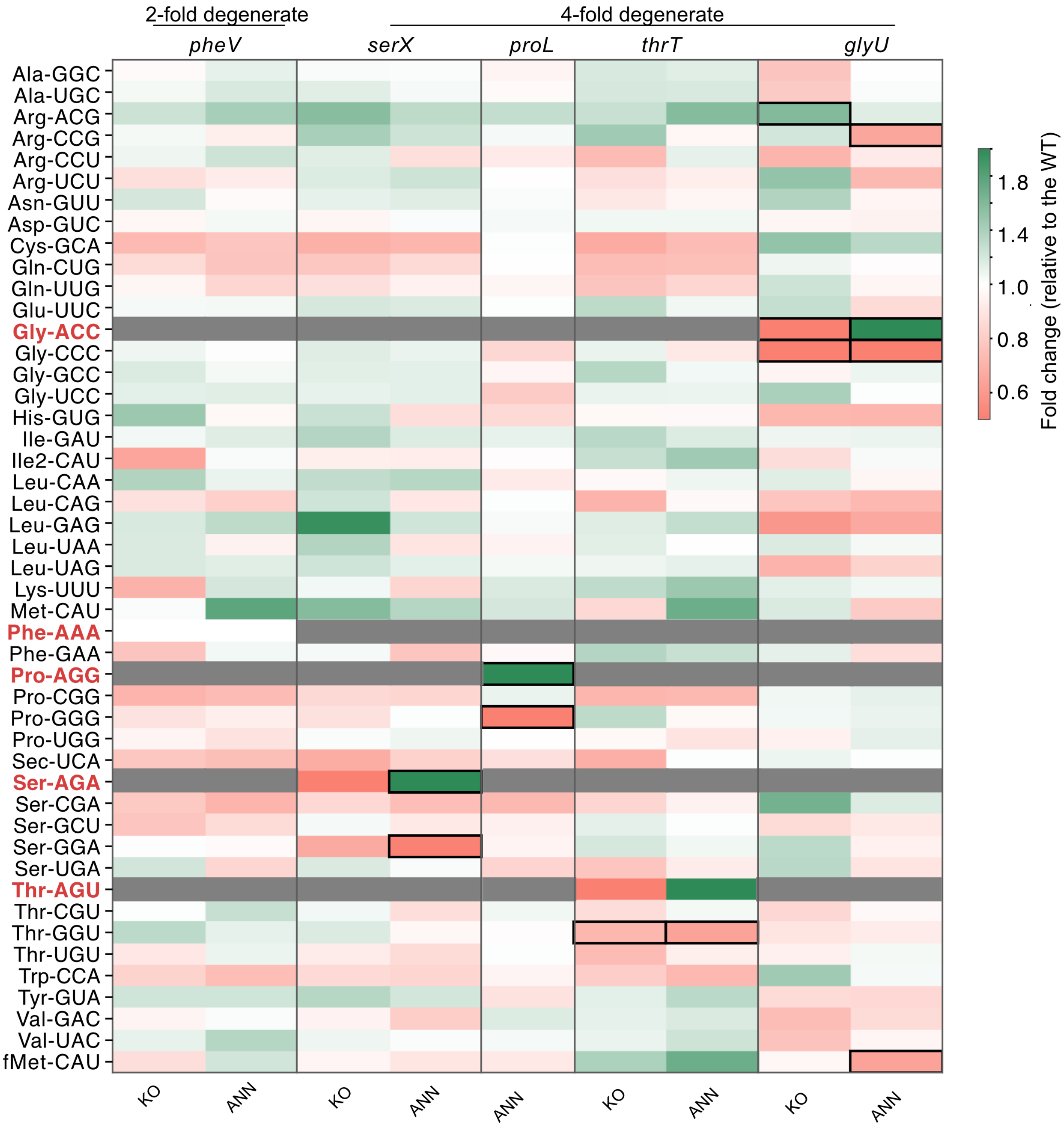

**Figure S4. Relative final OD and lag phase length of strains with a G34A substitution.** Final OD (**A**) and lag phase length **(B)** relative to the WT, for tRNA KO and strains with $tRNA_{ANN}$ expression from the genome (as measured in Fig. 2A). KO strains significantly different from WT and ANN strains significantly different from KO strains are indicated by thick borders.

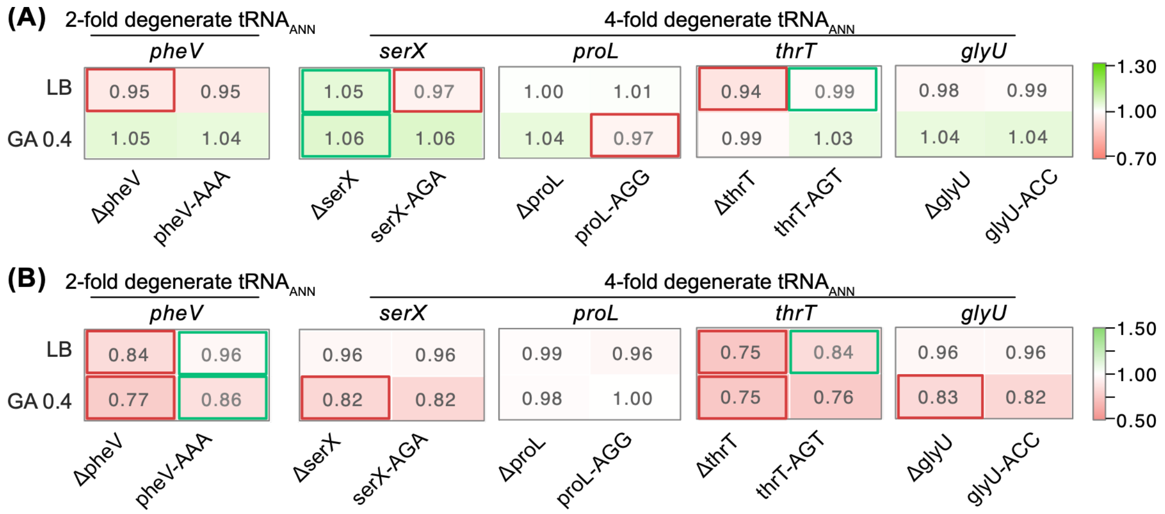

**Figure S5. Growth curves, lag phase length and final OD of strains lacking tRNA$_{BNN}$ complemented with respective tRNA$_{BNN}$ and tRNA$_{ANN}$. (A)** OD$_{600}$ v. time curves of tRNA knockouts (for *asnU*, *asnV*, *aspV*, *tyrV*, *proL*, *thrW* and *glyU* genes) complemented with their native tRNA$_{BNN}$ (control) or corresponding tRNA$_{ANN}$ variants from pUC19 vector. Each line shows mean OD$_{600}$ (n=4), with the shaded region denoting ±1 standard deviation. Growth media (labelled on the left) are arranged approximately in order of decreasing nutrient content: LB, GA 0.4 (M9 salts + 0.4% glucose + 0.4% cas amino acids), Glu 0.2 (M9 salts + 0.2% glucose), Gal 0.2 (M9 salts + 0.2% galactose), Gly 0.6 (M9 salts + 0.6% glycerol). Lag phase length **(B)** and final OD **(C)** relative to WT+pACDH (empty vector control), for KO+pACDH, KO+pACDH-tRNA$_{BNN}$ and KO+pACDH-tRNA$_{ANN}$ for a subset of genes (*asnU*, *asnV*, *aspV*, *tyrV*, *proL*, *thrW*, *glyU*). The KO+pACDH strains significantly different from WT+pACDH control and the complementation strains (KO+pACDH-tRNA$_{BNN}$ and KO+pACDH-tRNA$_{ANN}$) significantly different from the KO+pACDH are indicated by thick borders. Asterisks on the values of KO+pACDH-tRNA$_{ANN}$ indicate significant difference from KO+pACDH-tRNA$_{BNN}$. Significance is defined as to $p<0.05$ (Mann-Whitney U test). Cases where strains did not grow sufficiently well to calculate lag phase (e.g., tyrV-KO in Gly 0.6) are indicated by cells with a cross.

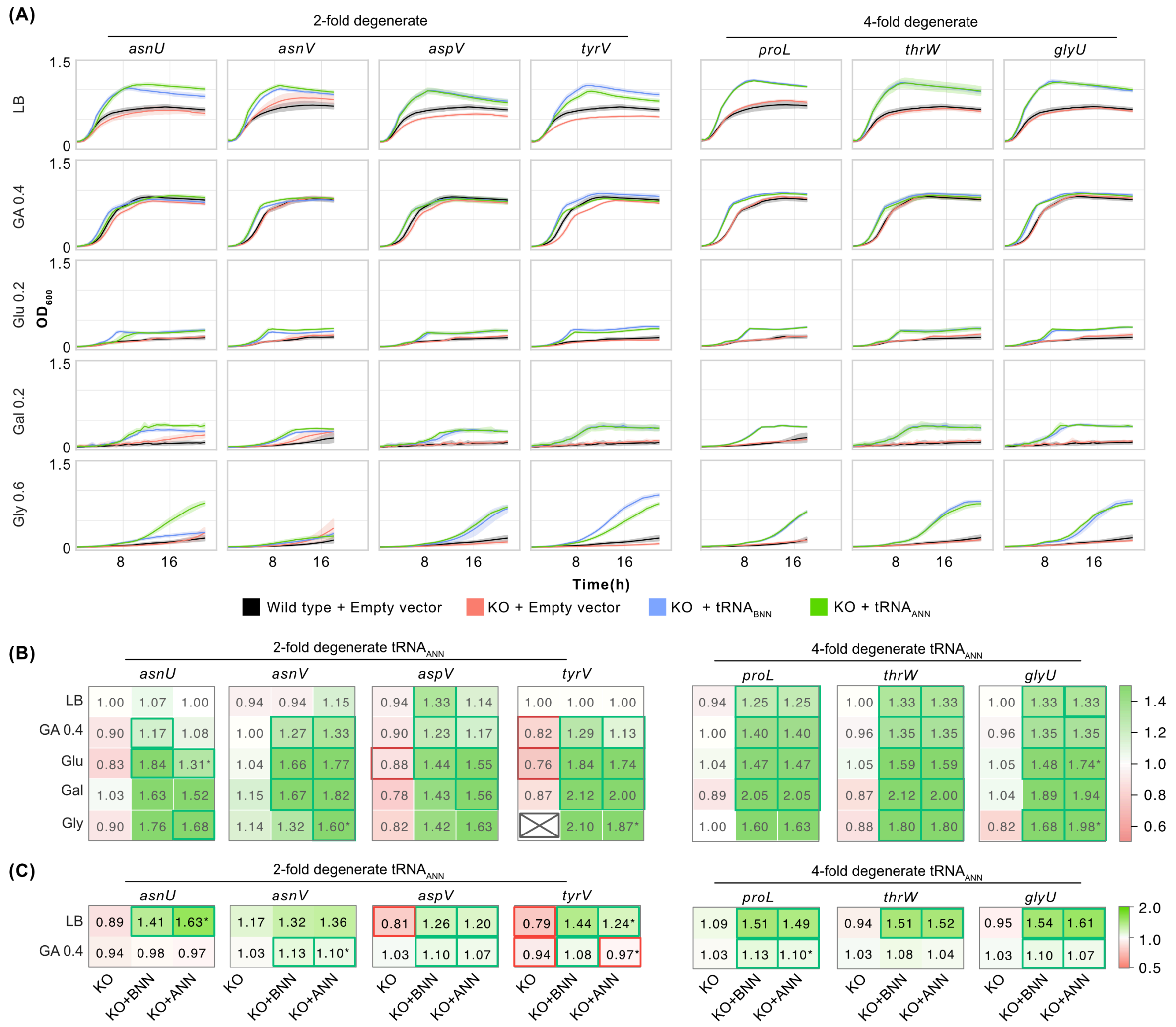

**Figure S6: Fitness effects of expressing $tRNA_{ANN}$ from a low copy number plasmid. (A)** $OD_{600}$ v. time curves for of the wild type (WT) expressing either $tRNA_{BNN}$ or corresponding $tRNA_{ANN}$ (each column; labelled on the top) from a low copy number vector across a concentration range of IPTG (each row; labelled on the left) in LB at 37°C. Each line shows mean $OD_{600}$ (n=4), with the shaded region denoting ±1 standard deviation. **(B–C)** Heat maps show the absolute growth rate **(B)** and the final $OD_{600}$ **(C)** from the $OD_{600}$ v. time curves shown in panel A. Cases where $tRNA_{ANN}$ were significantly different from $tRNA_{BNN}$ (Mann Whitney test, P value < 0.05; n=4) with an effect size higher than 5% are indicated by red ($tRNA_{ANN}$ < $tRNA_{BNN}$) or green ($tRNA_{ANN}$ > $tRNA_{BNN}$) thick borders. Statistically significant differences in effect sizes smaller than 5% (potentially within the range of noise resulting from detection limits and fitting of the growth equation) are indicated by thick grey borders. Heatmaps show growth rates **(D)** and final OD **(E)** of strains carrying $tRNA_{ANN}$ relative to that of their respective $tRNA_{BNN}$. Throughout the study, we compared fitness effects of overexpression of $tRNA_{ANN}$ with that of respective $tRNA_{BNN}$ to account for fitness effect stemming purely from overexpression. To calculate relative fitness parameters, we divided growth rate or OD of the strain expressing $tRNA_{ANN}$ by that of the strain expressing $tRNA_{BNN}$. Hence values of these parameters above 1 indicate a benefit (shown in green) and below 1 indicate a cost (shown in red) of $tRNA_{ANN}$ expression. tRNA genes were expressed on the plasmid pACDH and strains were grown in LB medium at 37°C across a gradient of inducer concentration (IPTG). Cases where $tRNA_{ANN}$ were significantly different from $tRNA_{BNN}$ (Mann Whitney test, $p<0.05$; N=4) with an effect size higher than 5% are further indicated by red or green thick borders. Statistically significant differences of effect sizes smaller than 5% (potentially within the range of noise resulting from detection limits and fitting of the growth equation) are indicated by thick grey borders.

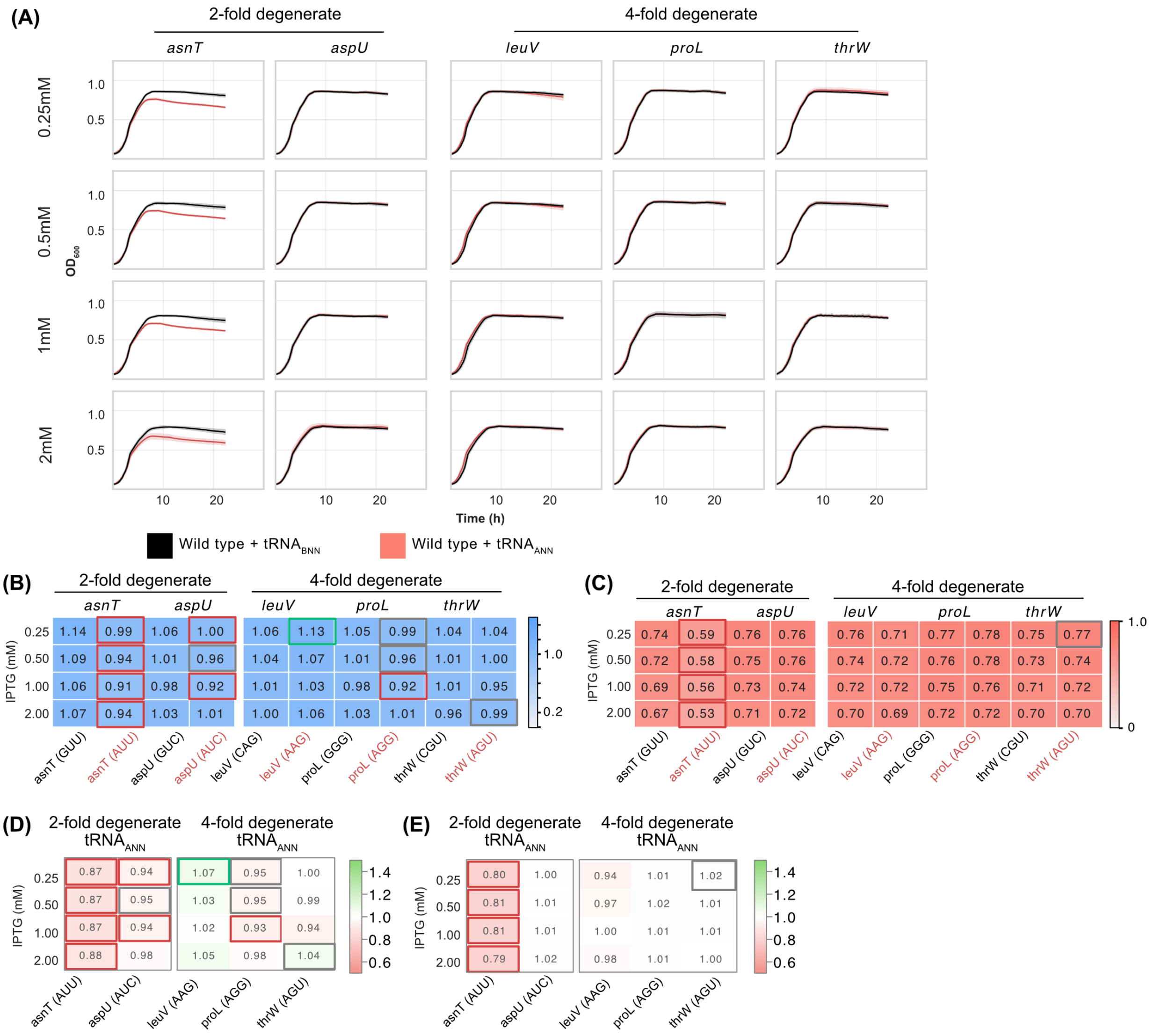
(A)
2-fold degenerate
4-fold degenerate
asnT
aspU
leuV
proL
thrW
0.25mM
0.5mM
1mM
2mM
$OD_{600}$
1.0
0.5
10
20
Time (h)
Wild type + $tRNA_{BNN}$
Wild type + $tRNA_{ANN}$
(B)
2-fold degenerate
4-fold degenerate
asnT aspU leuV proL thrW
IPTG (mM)
0.25 1.14 0.99 1.06 1.00 1.06 1.13 1.05 0.99 1.04 1.04
0.50 1.09 0.94 1.01 0.96 1.04 1.07 1.01 0.96 1.01 1.00
1.00 1.06 0.91 0.98 0.92 1.01 1.03 0.98 0.92 1.01 0.95
2.00 1.07 0.94 1.03 1.01 1.00 1.06 1.03 1.01 0.96 0.99
1.0
0.2
asnT (GUU) asnT (AUU) aspU (GUC) aspU (AUC) leuV (CAG) leuV (AAG) proL (GGG) proL (AGG) thrW (CGU) thrW (AGU)
(C)
2-fold degenerate
4-fold degenerate
asnT aspU leuV proL thrW
IPTG (mM)
0.25 0.74 0.59 0.76 0.76 0.76 0.71 0.77 0.78 0.75 0.77
0.50 0.72 0.58 0.75 0.76 0.74 0.72 0.76 0.78 0.73 0.74
1.00 0.69 0.56 0.73 0.74 0.72 0.72 0.75 0.76 0.71 0.72
2.00 0.67 0.53 0.71 0.72 0.70 0.69 0.72 0.72 0.70 0.70
1.0
0
asnT (GUU) asnT (AUU) aspU (GUC) aspU (AUC) leuV (CAG) leuV (AAG) proL (GGG) proL (AGG) thrW (CGU) thrW (AGU)
(D)
2-fold degenerate $tRNA_{ANN}$
4-fold degenerate $tRNA_{ANN}$
IPTG (mM)
0.25 0.87 0.94 1.07 0.95 1.00
0.50 0.87 0.95 1.03 0.95 0.99
1.00 0.87 0.94 1.02 0.93 0.94
2.00 0.88 0.98 1.05 0.98 1.04
1.4
1.2
1.0
0.8
0.6
asnT (AUU) aspU (AUC) leuV (AAG) proL (AGG) thrW (AGU)
(E)
2-fold degenerate $tRNA_{ANN}$
4-fold degenerate $tRNA_{ANN}$
IPTG (mM)
0.25 0.80 1.00 0.94 1.01 1.02
0.50 0.81 1.01 0.97 1.02 1.01
1.00 0.81 1.01 1.00 1.01 1.01
2.00 0.79 1.02 0.98 1.01 1.00
1.4
1.2
1.0
0.8
0.6
asnT (AUU) aspU (AUC) leuV (AAG) proL (AGG) thrW (AGU)

**Figure S7: Growth curves for strains expressing $tRNA_{ANN}$ from a high copy number plasmid.** $OD_{600}$ v. time curves for of the wild type (WT) expressing $tRNA_{BNN}$ and $tRNA_{ANN}$ (each column; labelled on the top) from a high copy vector across media (each row; labelled on the left) at 37°C, induced with 0.5 mM IPTG. Each line shows mean $OD_{600}$ (n=4), with the shaded region denoting ±1 standard deviation. **(A)** 2D codon $tRNA_{ANN}$ **(B)** 4D codon $tRNA_{ANN}$.

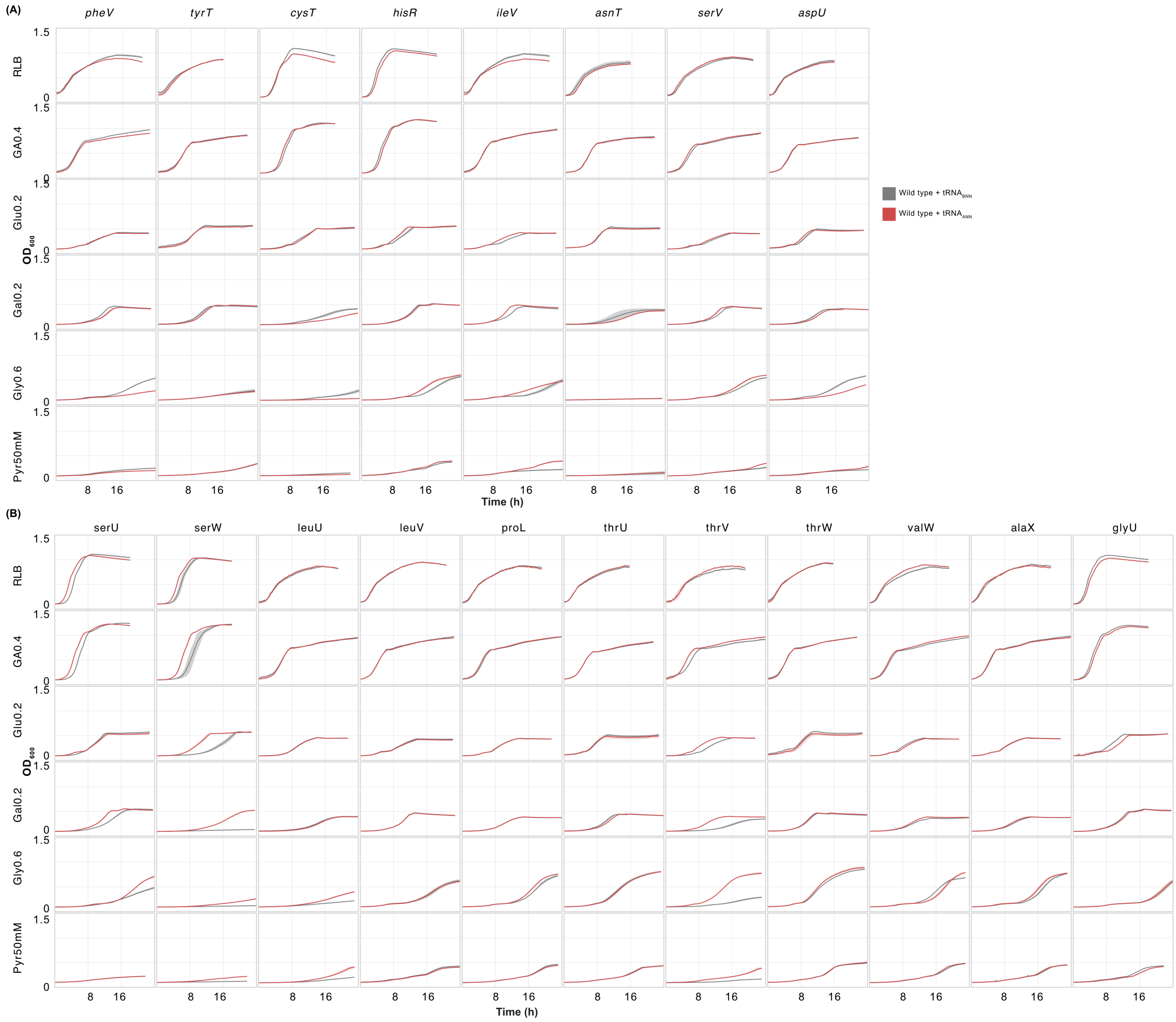

**Figure S8: Absolute growth parameters for strains expressing $tRNA_{ANN}$ from a high copy number plasmid.** Heat maps show the **(A)** absolute lag phase length, **(B)** growth rate and **(C)** final $OD_{600}$ estimated from the $OD_{600}$ v. time curves shown in Fig. S6. Cases where $tRNA_{ANN}$ were significantly different from $tRNA_{BNN}$ (Mann Whitney test, P value < 0.05; n=4) with an effect size > 5% are highlighted by red ($tRNA_{ANN}$ < $tRNA_{BNN}$) or green ($tRNA_{ANN}$ > $tRNA_{BNN}$) borders. Statistically significant differences in effect sizes < 5% (potentially within the range of noise resulting from detection limits and fitting of the growth equation) are indicated by thick grey borders. Empty cells in Fig. S7A indicate cases where strains did not grow sufficiently till the $OD_{600}$ threshold required to infer lag phase length.

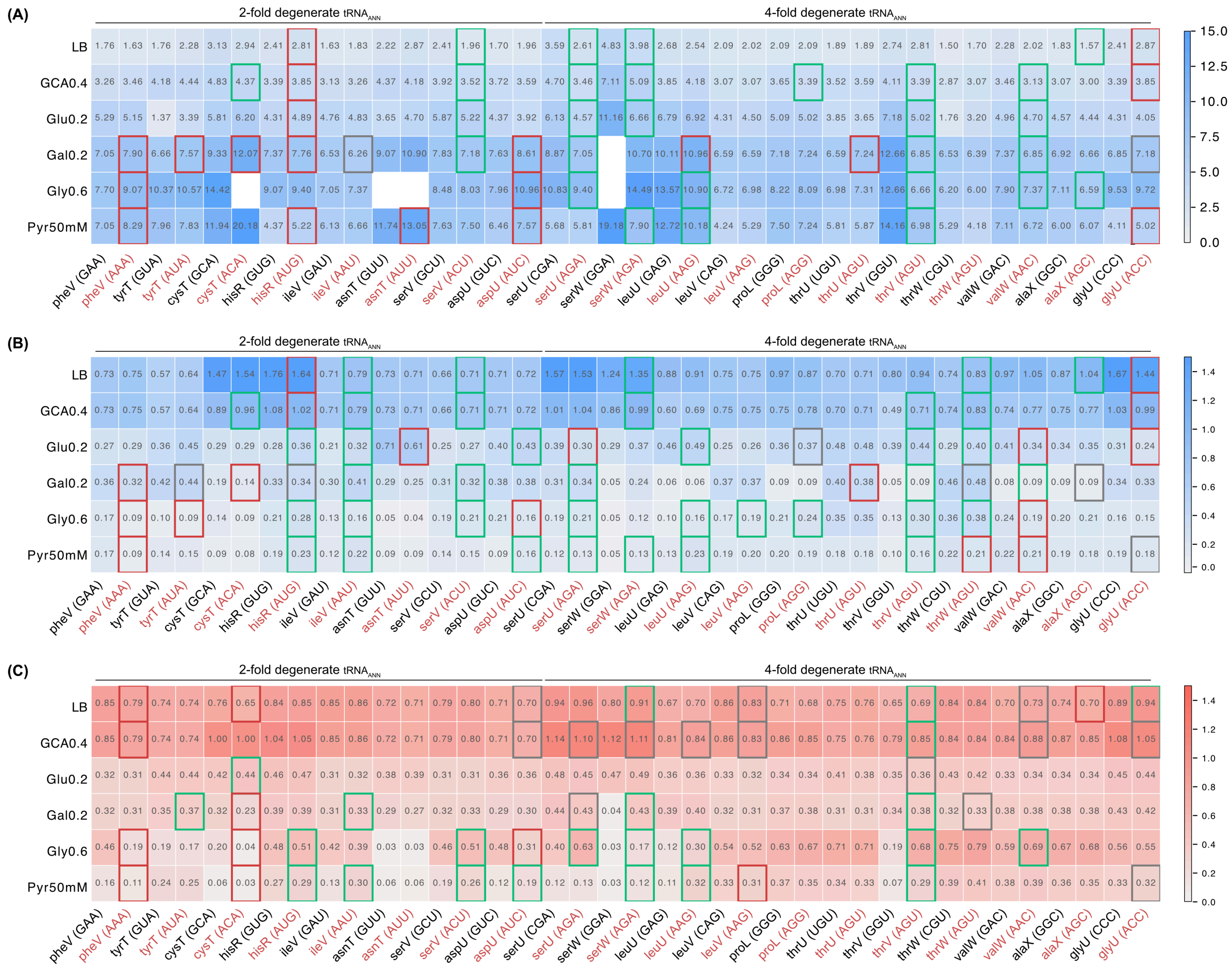

**Figure S9: Relative growth rate and yield for strains expressing $tRNA_{ANN}$ from a high copy number plasmid (A-B)** Heat maps show growth parameters of WT strains carrying $tRNA_{ANN}$, relative to those carrying the corresponding native $tRNA_{BNN}$ gene on the high copy plasmid pUC19 induced with 0.5mM IPTG at 37°C. Positive effects on growth ($tRNA_{ANN}$/ $tRNA_{BNN}$ > 1) are shown in green and negative impacts ($tRNA_{ANN}$/ $tRNA_{BNN}$ < 1) in red. Significant differences between $tRNA_{ANN}$ and the corresponding $tRNA_{BNN}$ (Mann-Whitney test, $p < 0.05$; N = 4) with effect size >5% are highlighted by a thick border. Statistically significant differences with effect size < 5% (potentially reflecting noise from detection limits and fitting of the growth equation) are indicated by thick grey borders. **(A)** relative growth rate and **(B)** relative final OD are shown across different media, with values of each parameter given in each cell. The first two rows include nutrient-rich media and the next four rows include nutrient-poor media. In some cases, data for one or more replicates from either the $tRNA_{ANN}$ strain or the respective $tRNA_{BNN}$ variant did not fit the exponential growth equation whereas the other variant grew; e.g., Ser-$tRNA_{GGA}$ in Gal 0.2 medium did not grow, unlike Ser-$tRNA_{AGA}$ (raw $OD_{600}$ vs. time curves are shown in Fig. S7). Such cases yielded infinite or infinitesimal values for relative growth rate, so the corresponding cells in the heatmaps are empty; but the fill colours and outlines indicate the qualitative direction of relative growth (green if only $tRNA_{ANN}$ grew and red if only $tRNA_{BNN}$ grew). The absolute values of growth parameters are shown in Fig. S8 and summarised in the source data file for Fig. S8. **(C)** Heatmaps show qualitative fitness effects of each $tRNA_{ANN}$ on the three growth parameters across media (positive, negative and neutral effects shown in green, red and white, respectively).

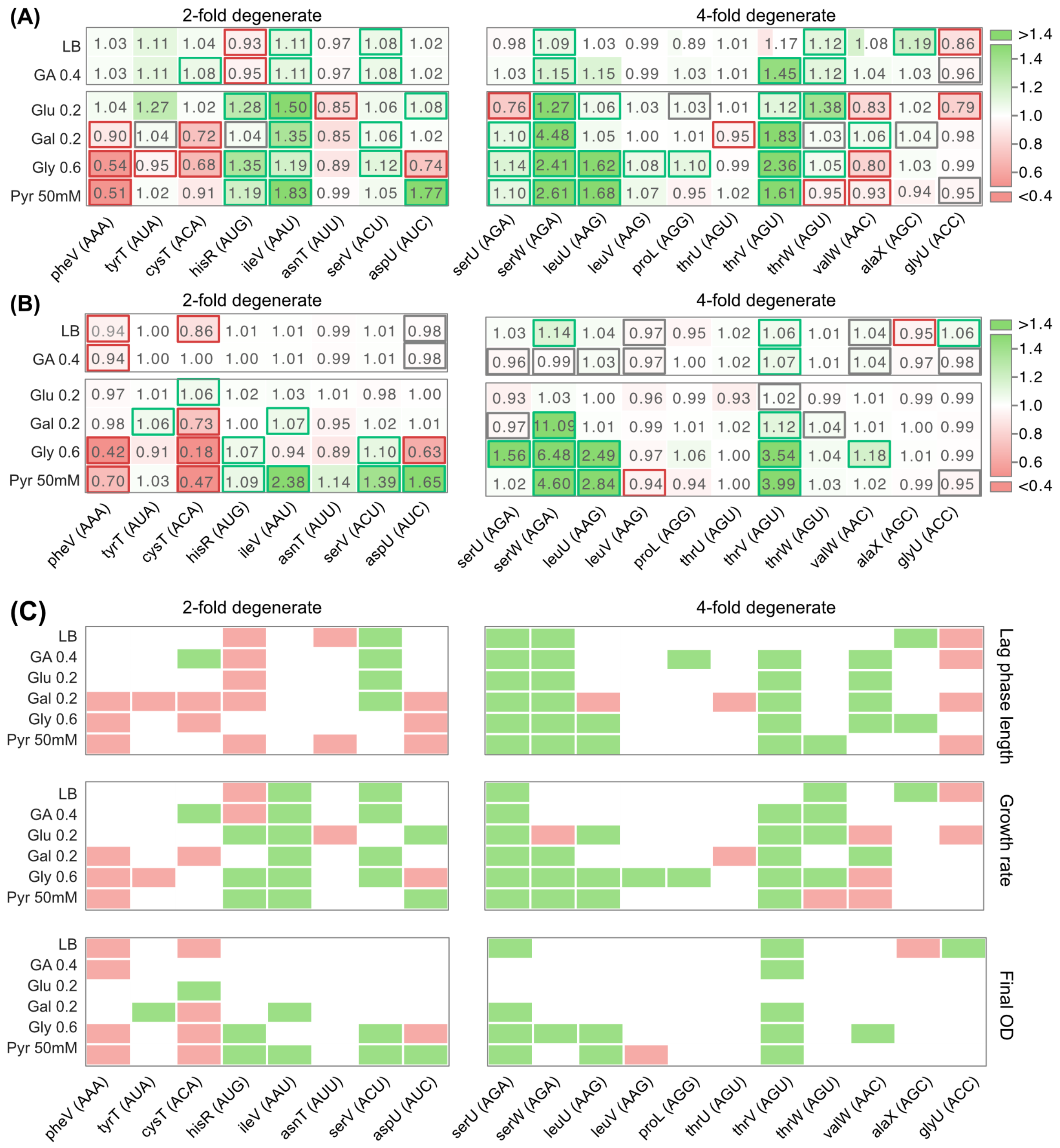

(A)
(B)
(C)
2-fold degenerate
4-fold degenerate
LB
GA 0.4
Glu 0.2
Gal 0.2
Gly 0.6
Pyr 50mM
pheV (AAA)
tyrT (AUA)
cysT (ACA)
hisR (AUG)
ileV (AAU)
asnT (AUU)
serV (ACU)
aspU (AUC)
serU (AGA)
serW (AGA)
leuU (AAG)
leuV (AAG)
proL (AGG)
thrU (AGU)
thrV (AGU)
thrW (AGU)
valW (AAC)
alaX (AGC)
glyU (ACC)
>1.4
1.4
1.2
1.0
0.8
0.6
<0.4
Lag phase length
Growth rate
Final OD

**Figure S10: Fitness effects of $tRNA_{ANN}$ and estimated mistranslation likelihood in strains expressing split-codon $tRNA_{ANN}$. (A-B)** Schematics illustrate decoding in examples of 2D and 4D codon boxes under Watson-Crick (WC; indicated by black lines), wobble (indicated by dotted black lines) and superwobble (indicated by dotted orange lines) in the wild type (WT) and after B34A substitutions in distinct $tRNA_{BNN}$. In each case, mRNA codons are written from 5'-to-3' at the bottom, and complementary anticodons written from 3'-to-5' on top, with flanking lines and attached single letter codes indicating the tRNA backbone and amino acid after charging, respectively. The $34^{th}$ base is shown in bold and anticodon mutations leading to A34 are marked in red. Predicted mistranslation is indicated by red arrows. **(A)** A codon box split between phenylalanine and leucine, with two codons each decoded by Phe-tRNA and Leu-tRNA in the WT. A G34A substitution would generate Phe-$tRNA_{AAA}$ charged with phenylalanine, which under WC pairing reads the phenylalanine codon UUU but can decode the leucine codons UUA and UUG through superwobbling, resulting in mistranslation. Similarly, U34A or C34A substitutions would result in $tRNA_{AAA}$ charged with leucine, resulting in phenylalanine-to-leucine mistranslation. Superwobbling resulting in mistranslation is indicated by red arrows and is expected to be harmful. **(B)** The leucine 4D codon box, where all four codons are decoded by $tRNA_{GAG}$, $tRNA_{UAG}$ and $tRNA_{CAG}$ isoacceptors. A B34A substitution in any of these isoacceptors should generate Leu-$tRNA_{AAG}$ charged with leucine. Under canonical base pairing, Leu-$tRNA_{AAG}$ decodes the CUU codon and under superwobbling it decodes CUC, CUA and CUG. In either case, the leucine charged Leu-$tRNA_{AAG}$ decodes leucine codons, and does not result in mistranslation. Hence, in $tRNA_{BNN}$ from 4D codon boxes, B34A mutations are unlikely to mistranslate. Superwobbling by $tRNA_{ANN}$ within 4D codon boxes is indicated by green arrows and is expected to be tolerated or beneficial. Such superwobbling is indicated only for a $tRNA_{ANN}$ resulting from G34A substitution, but it is equally applicable for any B34A substitution within a 4D codon box. **(C)** Illustration of how we estimated mistranslation probability for 2D $tRNA_{ANN}$, assuming that $tRNA_{ANN}$ will outcompete $tRNA_{BNN}$ and decode the relevant codon box. The probability of amino acid misincorporation ('mistranslation') is estimated as the proportion of non-cognate (U- and C-ending) codons in a given codon box, as illustrated for the Phe-Leu split codon (2D) box (paratheses show the genome-wide usage of each codon). **(D)** Relationship between the fitness effect index and mistranslation probability for the eight 2D codon $tRNA_{ANN}$. The dotted line indicates a linear regression fit with 95% CI. To calculate fitness effect index, we assigned qualitative fitness scores (+1, 0, and –1 to positive, neutral, and negative effects, respectively) to each strain-media from the figure S9C. The sum of these scores, across media and across parameters, was defined as fitness effect index.

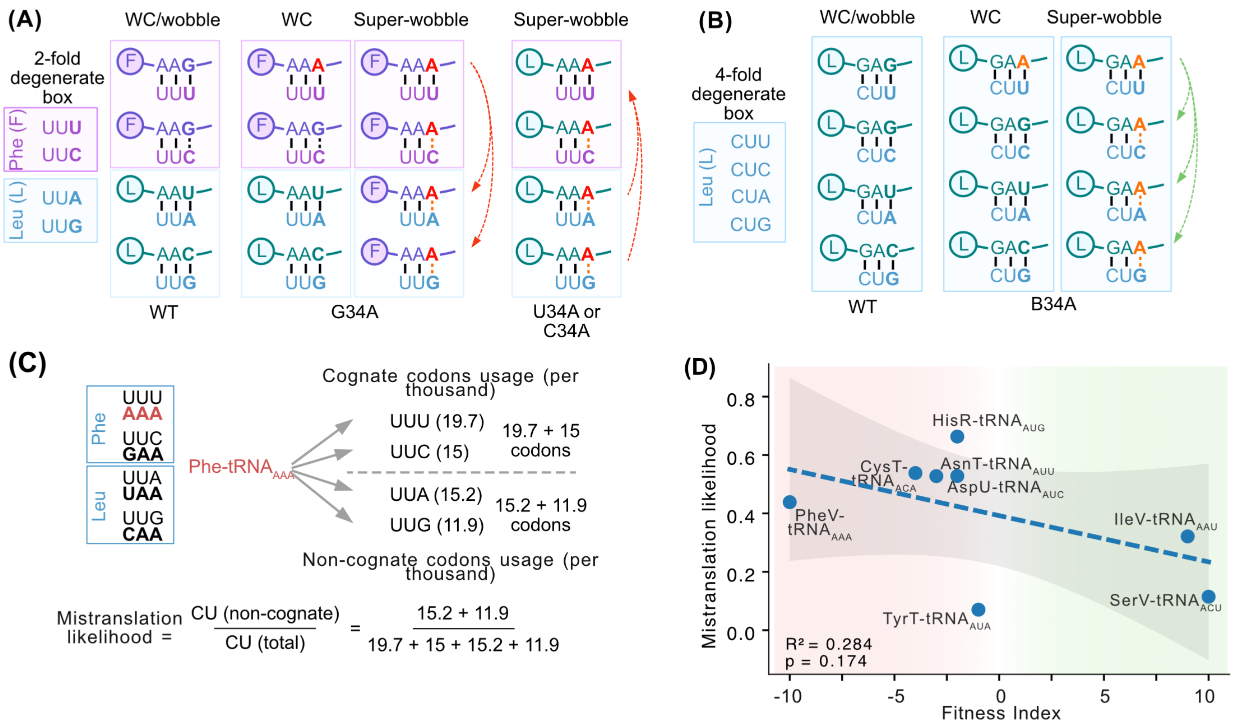

(A)
WC/wobble
WC
Super-wobble
Super-wobble
2-fold degenerate box
Phe (F)
UUU
UUC
Leu (L)
UUA
UUG
WT
G34A
U34A or C34A
(B)
WC/wobble
WC
Super-wobble
4-fold degenerate box
Leu (L)
CUU
CUC
CUA
CUG
WT
B34A
(C)
Phe
UUU
AAA
UUC
GAA
Leu
UUA
UAA
UUG
CAA
Phe-tRNA AAA
Cognate codons usage (per thousand)
UUU (19.7)
UUC (15)
19.7 + 15 codons
UUA (15.2)
UUG (11.9)
15.2 + 11.9 codons
Non-cognate codons usage (per thousand)
Mistranslation likelihood = CU (non-cognate) / CU (total) = 15.2 + 11.9 / 19.7 + 15 + 15.2 + 11.9
(D)
Mistranslation likelihood
Fitness Index
HisR-tRNA AUG
CysT-tRNA ACA
AsnT-tRNA AUU
AspU-tRNA AUC
PheV-tRNA AAA
IleV-tRNA AAU
SerV-tRNA ACU
TyrT-tRNA AUA
R² = 0.284
p = 0.174

**Figure S11: Occurrence of tRNA$_{ANN}$ (other than Arg-tRNA$_{ACG}$) across bacteria.** Presence of tRNA$_{ANN}$ from 2D (red cells) and 4D (green cells) codon boxes, as predicted by GtRNAdb, across 555 bacterial genomes grouped by phylum (see the source data for individual species and full taxonomy). The last column shows presence of a TadA homolog (green cells). White cells indicate absence of the tRNA/*tadA* gene; grey cells in the tadA column indicate 16 species where amino acid CDS could not be retrieved from NCBI based on the taxonomic ID provided in GtRNAdb.

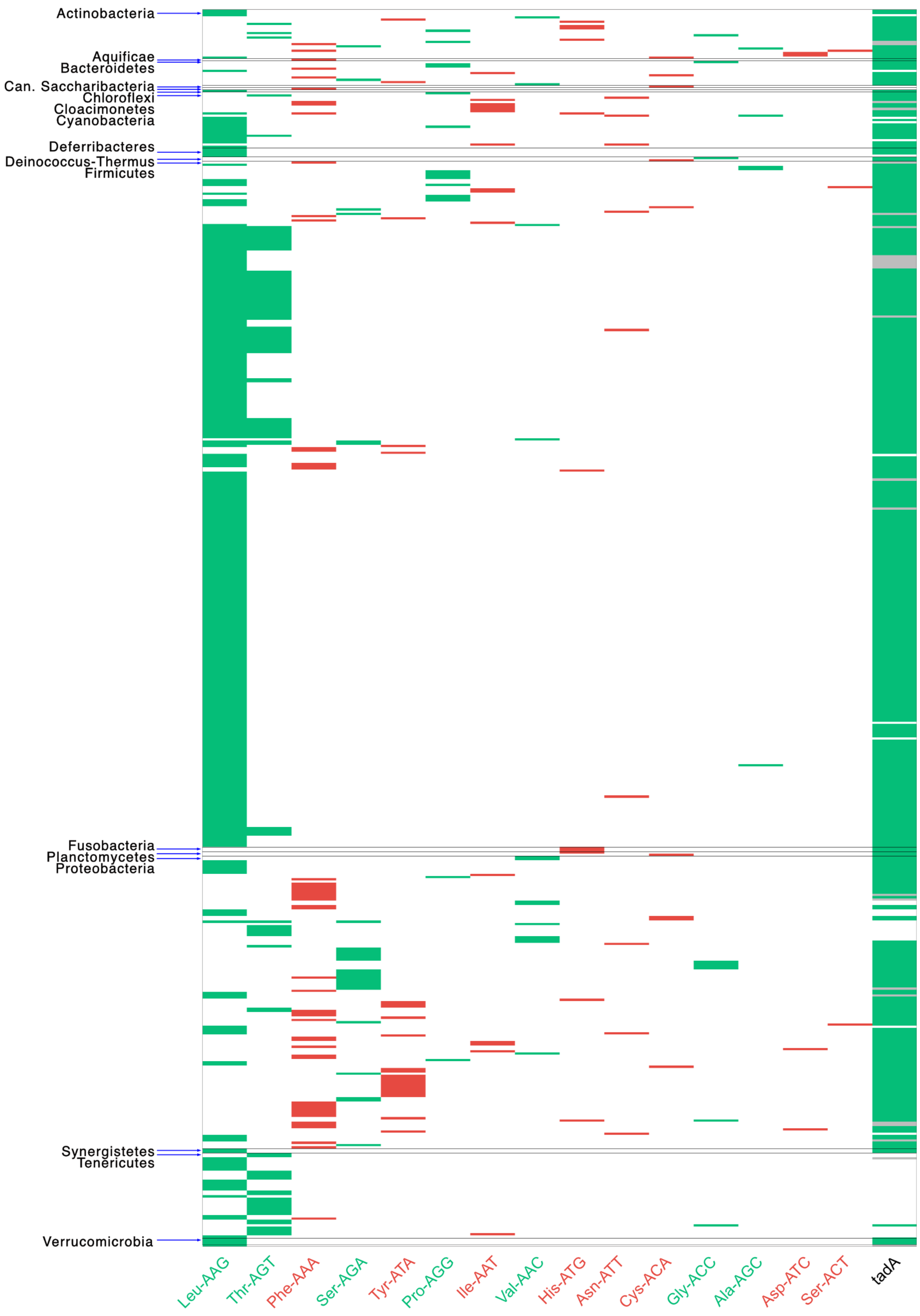

**Figure S12: Co-occurrence of $tRNA_{ANN}$ (other than Arg-$tRNA_{ACG}$) with their isoacceptors.** Out of all species with a given $tRNA_{ANN}$ (other than Arg-$tRNA_{ACG}$), the fraction of species that also encode isoacceptor $tRNA_{TNN}$, $tRNA_{CNN}$ and $tRNA_{GNN}$.

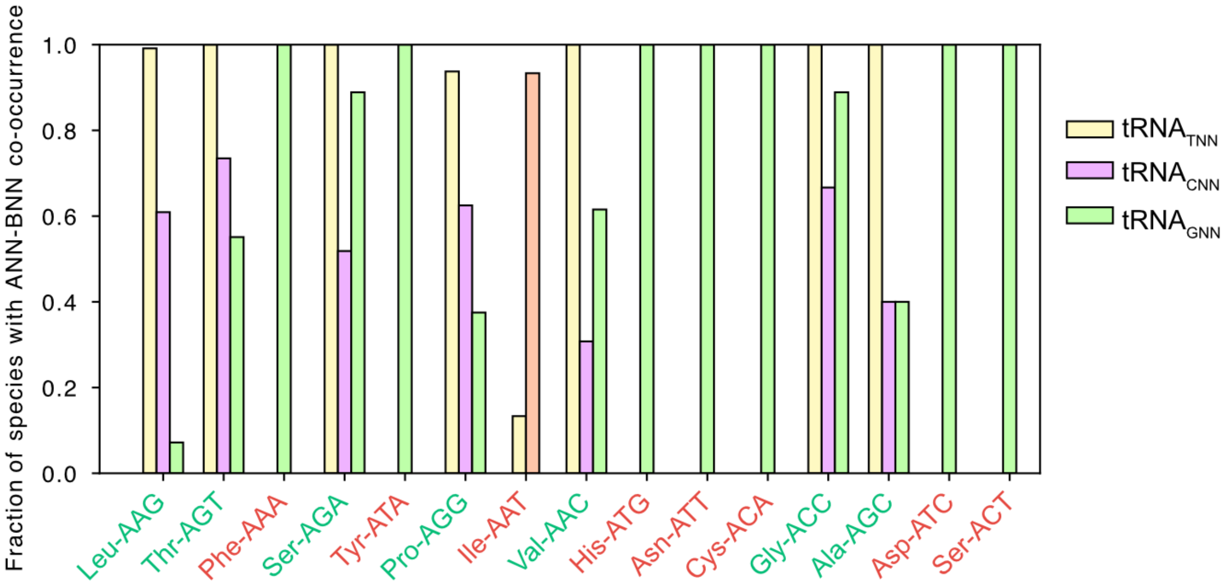

**Figure S13: Fate of an A34 allele with distinct selection coefficients.** Wright-Fisher simulations of the frequency of a single A34 allele (with varying selection coefficient s) arising in a haploid population of size $N_e = 10^9$. At each generation t, the frequency of the A34 allele ($P_{A34}$) was calculated as $p_{t-1}(1+s) / (1 + sp_{t-1})$, where $p_{t-1}$=frequency of A34 allele in the previous generation. The number of A34 alleles in generation t+1 ($X_{t+1}$) was sampled from a binomial distribution ($N_e$, $P_{A34}$), given the frequency $p_{t+1}=X_{t+1}/N_e$. This was iterated until $p_{t+1}=1$ (fixation) or $p_{t+1}=0$ (extinction), and the number of generations (t) to reach each outcome was recorded. Each panel shows the outcomes of 1000 such simulations with varying s (indicated on top of each panel), with the percentage of simulations in which the A34 allele was fixed or went extinct and the average number of generations taken for each outcome.

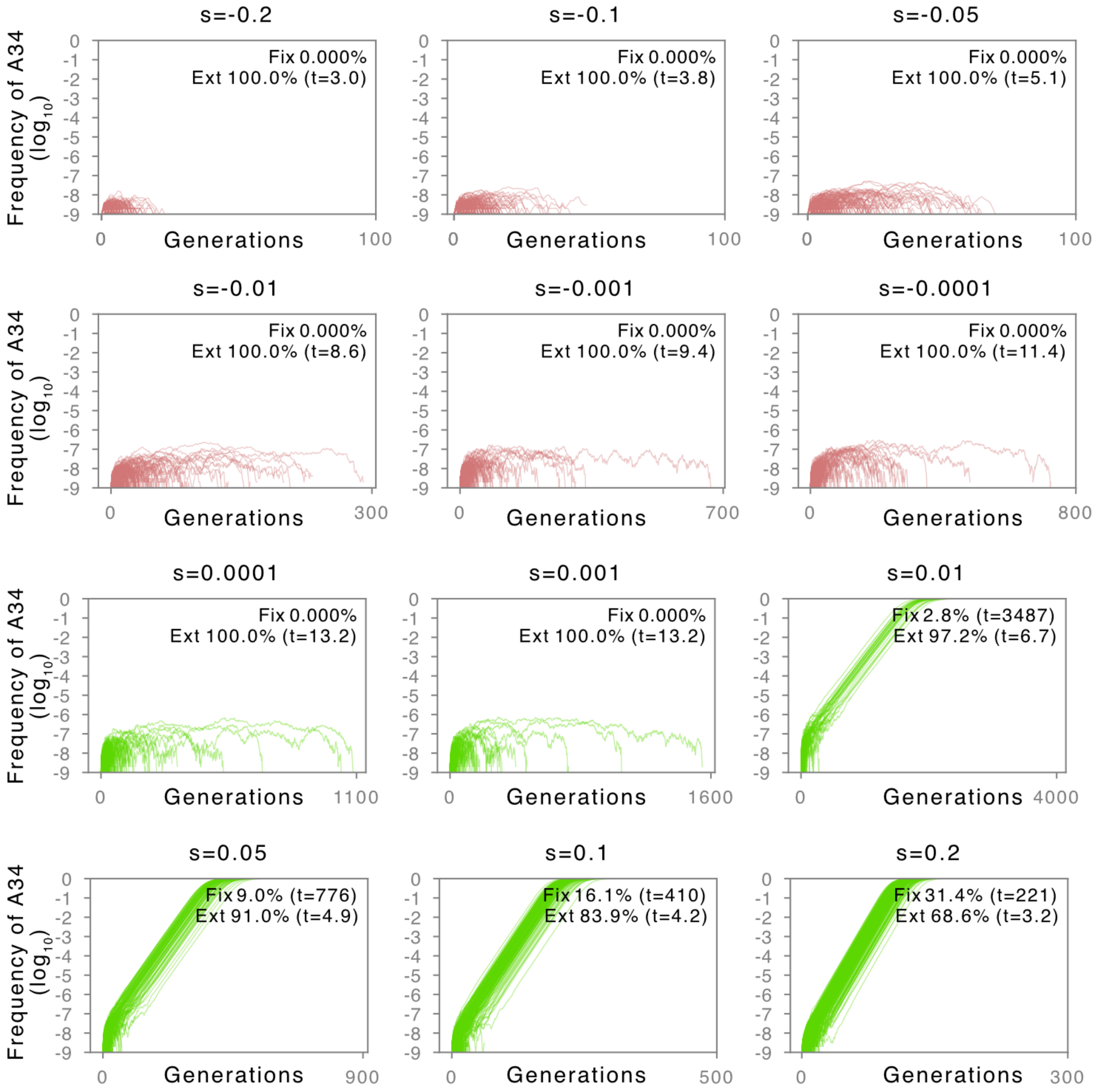
